# Regenerative clustering of Enteroblasts in the *Drosophila* midgut revealed by a morphometric analysis

**DOI:** 10.1101/2024.03.10.584282

**Authors:** Fionna Zhu, Michael J. Murray

## Abstract

Enteroblasts (EBs) are the cells responsible for the maintenance of the epithelium that lines the adult midgut in *Drosophila*. In response to cell death and damage, EBs undergo a Mesenchymal-Epithelial-Transition (MET) as they incorporate into the epithelium and differentiate into enterocytes (ECs). The morphogenetic mechanisms driving this MET process are not well understood. To improve phenotypic analysis of EBs, we established an analysis pipeline that uses machine learning segmentation to produce a reliable and automated quantification of cellular morphology and spatial distributions. EB morphology and fate is visualised using the Repressible Dual Differential stability cell Marker (ReDDM) approach. We show that wildtype EB cells exhibit a bimodal distribution pattern in which midguts fall into two categories: “quiescent” guts, in which EBs are evenly spaced out and newly formed ECs are uncommon, and “regenerative” guts, in which EBs are clustered and new ECs are prevalent. Using this system we first show that RNAi knockdown of Septate Junction proteins disrupts normal EB morphology and spatial distribution. With time-lapse imaging, we have also established that EBs are motile in nature, and when artificial tissue damage was introduced, exhibited increased cytoplasmic movements, and formed distinct clusters. We then demonstrate the utility of our pipeline in a small candidate screen for genes that might mediate EB clustering. Based on our results, we propose a working model that links the dynamic behaviour of EBs with midgut regeneration.

## Introduction

Homeostatic mechanisms are essential for maintaining the proper function of epithelial tissues in the adult body. This process of replenishing old or damaged cells by less differentiated progenitors to re-establish homeostasis is known as cell turnover (Pellettieri & Alvarado, 2007). The rate and manner of homeostatic cell turnover varies across different epithelial tissues, ranging from constant renewal (Flier and Clevers, 2009), to hormone-dependent renewal (Gargett et al., 2008), and finally delayed turnover in slow-cycling epithelia (Rawlins and Hogan, 2006). In the last category, stem cells reside in a basal compartment and supply progenitor cells that replace epithelial cells as needed.

Here we utilised a simple epithelium - the adult *Drosophila* midgut to investigate how epithelial homeostasis is maintained by “on-demand” progenitor differentiation. The *Drosophila melanogaster* midgut is functionally equivalent to the mammalian small intestine and has emerged as a powerful model system to study intestinal dysfunction (Biteau et al., 2011). Structurally, the *Drosophila* midgut is a monolayered epithelium comprised of 4 cell types: the absorptive Enterocytes (ECs) and the secretory Enteroendocrine cells (EEs) that form the epithelium, and the basally positioned intestinal stem cells (ISCs) and intermediate progenitor Enteroblasts (EBs) (Micchelli and Perrimon, 2006; Ohlstein and Spradling, 2006). ISC cells undergo symmetric and asymmetric division giving rise to ISCs, pre-EEs, or EBs. ECs arise from EBs following a Mesenchymal-Epithelial Transition (MET) governed by Notch signalling, and EEs are generated through a designated set of progenitors known as pre-EEs (Biteau and Jasper, 2014; He et al., 2018; Zeng and Hou, 2015).

Following damage, midgut regeneration is almost exclusively dependent on rapid ISC proliferation and EB differentiation to restore lost cells (Amcheslavsky et al., 2009; Buchon et al., 2009b; Jiang et al., 2009, 2016; Tian et al., 2017), though occasionally the regenerative ISC pool can be derived through EB divisions (Tian et al., 2022). The balance between cell proliferation and differentiation must be carefully regulated, as the molecular mechanisms underpinning tissue regeneration and tumourigenesis are very similar (Arwert et al., 2012). For this reason, there has been an abundance of studies on the signalling mechanisms controlling stem cell behaviour during epithelial homeostasis (Zhang and Edgar, 2021). However, the mechanical aspects of the MET event that occurs during EB differentiation into EC are less well known. Recently, one aspect of the EB-to-EC differentiation mechanics was described in detail. Work by two independent labs highlighted the importance of luminal surface formation in proper EB differentiation, which requires the coordination between cell identity transition and the intercalation into a pre-existing epithelium. This is accomplished through the formation of a pre-assembled apical compartment (PAC) (Chen and St Johnston, 2022; Galenza et al., 2023).

MET is a cellular process that describes the phenotypic transition from a migratory mesenchymal cell state to a stationary epithelial cell state (Thiery et al., 2009). The essence of MET is the transition from front-back polarity to apical-basal polarity, and the establishment of functionally distinct domains and specialised cell-cell junctions (Bryant and Mostov, 2008). MET is involved in development, tissue homeostasis and disease progression (Chaffer et al., 2007). Three strategies of MET are used by mesenchymal cells to transition into polarised epithelial cells: single cell MET, propagative MET, and collective MET (Kim et al., 2017). Amongst the three, single cell MET most accurately describes the EB-to-EC MET event which occurs in an established tissue, whereby a mesenchymal cell integrates itself into the existing epithelium.

Previous studies have revealed a number of master regulators of EB MET: they are the transcription factors *escargot (esg)*, *Sox21a*, *zfh2*, and *microRNA-8* (*miR-8*). EB differentiation can be triggered by Zfh2, which was thought to acts upstream of Sox21a and miR-8 (Rojas Villa et al., 2019). miR-8 drives EB MET by repressing the transcription factor *esg*, which functions to maintain the cell’s mesenchymal state (Antonello et al., 2015b, 2015a; Korzelius et al., 2014; Loza-Coll et al., 2014; Nègre et al., 2011). Multiple signalling pathways have also been shown to be involved in the EB-to-EC differentiation: Target of Rapamycin (TOR) signalling, insulin signalling and EGFR signalling (Choi et al., 2011; Kapuria et al., 2012; Xiang et al., 2017). Events that increase the need for regeneration such as mechanical injury or bacterial infection result in an increase in the number of miR-8 positive EBs (Antonello et al., 2015a), though the precise mechanism underlying how EBs perceive such damages remained largely unelucidated.

While the regulation of EB production and differentiation have also been extensively studied, the analyses often focus on the genes expressed by these cells, and not their morphological and positional characteristics. Studies that do describe the morphological transformations of EB cells tend to be qualitative in nature, with only a handful taking a more quantitative approach. In one study, manipulating *Zfh2* levels in both ISCs and EBs led to differential effects on cell area and circularity (Rojas Villa et al., 2019). In another study, knock down of *dronc* affected EB shape and size, though only size was quantified (Lindblad et al., 2021). These studies are evidence that gene effects on cell status can be reflected through their morphological changes. As the morphological features of EB cells are a key aspect of EB activation and differentiation, being able to measure and quantify them would allow for more robust comparisons of different genotypes and treatments.

Here we describe an analysis pipeline for a comprehensive and quantitative measurement of EB morphology and spatial distribution data in the adult *Drosophila* midgut. This method uses the Repressible Dual Differential stability cell Marker (ReDDM) system to label EBs and newly formed ECs (Antonello et al., 2015a) and integrates machine-learning based image segmentation (ilastik) (Berg et al., 2019), morphological shape analysis (ImageJ), and spatial statistics (Andrey et al., 2010). With this pipeline, we characterised wildtype EBs and uncovered a previously unreported bimodality of spatial distribution in which midguts tended to be either “quiescent” or “regenerative”: quiescent guts had EBs evenly spread out and very few newly formed ECs, whilst regenerative guts, had regions that we call “regenerative clusters” in which EBs and new ECs were clustered together. Using the pipeline we analysed genotypes in which key categories of EB genes are knocked down and found that loss of Septate Junction proteins in EBs significantly disrupts normal cell morphology, suggesting that these molecules are not only crucial for differentiation and junction formation, but also in the maintenance of the undifferentiated states of EB cells. This method also allowed us to examine EB behaviour during regeneration when disruptive chemicals (i.e. DSS) were fed, through fixed tissue immunostaining and ex vivo culture live imaging. Finally, we used our pipeline to conduct an RNAi screen for genes involved in regeneration, and uncovered several mechanosensors that are implicated in Regenerative cluster formation. Together, we propose a working model in which EB cells sense nearby damage sites through mechanosensors or surface receptors, actively migrate towards the damage site and respond to damage in a collective fashion via the formation of Regenerative clusters. Together, our results demonstrate the wealth of image data which is currently underutilised and its capacity to improve our ability to compare and understand gene function and cellular behaviour.

## Materials and Methods

### Fly husbandry

Flies were raised and maintained on standard molasses fly food. All crosses were conducted in an 18°C incubator unless otherwise specified. Experimental virgin females were kept at 18°C for 3 days before transferring to a 29°C incubator to induce Gal4 expression. 3-day old virgin females were crossed to *w^1118^* males before shifting to 29°C. All animals used for analyses were dissected at 10-days of age. Unless otherwise stated, for all ReDDM experiments, flies were turned onto new food every 5 days. For MET regenerative cluster experiments, flies were turned onto new food at frequencies as described in the main text.

### Drosophila stocks

Strains of *Drosophila melanogaster* were obtained from Bloomington Stock Centre (http://flystocks.bio.indiana.edu) or Vienna *Drosophila* Resource Centre (https://stockcenter.vdrc.at) unless otherwise indicated. Genetic symbols are described in Flybase (http://flybase.org). The *w^1118^* strain was used as wildtype control for phenotypic analysis. The following stocks were used in this study:

*Su(H)GBE-Gal4, UAS-mCD8-GFP/CyO; gal80^ts^, UAS-H2B-RFP/TM6B* (this work, made from *esg-Gal4, UAS-mCD8-GFP/CyO; gal80^ts^, UAS-H2B-RFP/TM2* (a gift from Dr. Maria Dominguez)). *UAS-Ssk.IR^V105193^, UAS-mesh.IR^V040940^, UAS-Itgbn.IR^BL28601^, UAS-Rac1.IR^BL34910^, UAS-slik.IR^BL35179^, UAS-Piezo.IR^VDRC2796^, UAS-nompC.IR^BL31512^, UAS-pain.IR^BL31511^, UAS-nan.IR^BL53312^, UAS-wrtw.IR^BL51503^, UAS-trpm.IR^BL57871^, UAS-ppk.IR^BL29571^, UAS-ppk29.IR^BL27241^, UAS-sand.IR^BL25853^, UAS-arm.IR^BL31304^, UAS-fz2.IR^BL31312^, UAS-moody.IR^BL66326^, UAS-pax.IR^BL28695^, UAS-FAK.IR^BL33617^, UAS-cac.IR^BL27244^, UAS-yki.IR^BL31965^, UAS-mys.IR^BL33642^, UAS-mew.IR^BL44553^, UAS-fra.IR^BL31469^, UAS-dome.IR^BL34618^, UAS-pvr.IR^BL37520^, UAS-dscam1.IR^BL38945^, UAS-dscam2.IR^BL51839^, UAS-dscam4.IR^BL51508^, UAS-Ed.IR^BL38243^, UAS-Egfr.IR^V043268^, UAS-btl.IR^BL55870^, Su(H)GBE-lacZ (a gift from Dr. Gary Hime); Su(H)GBE-lacZ; pUAST-FRT-Stop-FRT/CyO (generated from pUAST-FRT-Stop-FRT-rpr/CyO; byn >Gal4, Tub >Gal80/TM6; a gift from Dr. Louise Cheng. Cohen et al., (2018)), hsFLP, UAS-mCD8-GFP; GMR45D10-GAL4/TM6B (generated from hsFLP, UAS-mCD8-GFP/FM7; tub-GAL4, FRT82B tubP-GAL80/TM6B (a gift from Dr. Leonie Quinn), and GMR45D10-GAL4^BL45323^)*.

### *Drosophila* adult midgut dissection

Only mated adult females were used in this work. Flies were anesthetised on a CO2 block prior to dissection. A pair of slightly blunt tweezers were used to hold the appendages of the animal tightly, very close to its body. The animal was then submerged in 2mL of 1x PBS solution inside the cavity block. A pair of sharp tweezers were used to remove both wings. The back cuticle was removed to eliminate air bubbles. The head of the animal was removed, followed by opening of the thorax and gently lifting out the anterior portion of the midgut. The proventriculus and the crop were left intact and attached to the rest of the midgut. The abdomen was opened, and the midgut was removed by carefully severing it from the hindgut and any other tissue attached (e.g. ovaries and fat body). This process was conducted in under 5 minutes. Dissected midguts were used for fixed imaging as well as ex vivo cultured time-lapse imaging.

### *Drosophila* adult midgut fixation and immunostaining

Dissected midguts were fixed in 4% formaldehyde (diluted from 16% EM-grade formaldehyde, ProSciTech) in 1x PBS for 45 minutes in room temperature on a gyratory mixer. The 16% formaldehyde ampoules were aliquoted into 1ml aliquots and kept at 4°C prior to usage. Fixed midguts were rinsed and washed with PBST (PBS, 0.1% Triton-X) for 30 minutes prior to immunostaining. All antibodies were buffered in PBST. Midguts were incubated in primary antibodies overnight at 4°C and secondary antibodies for 2 hours at room temperature on an orbital mixer. The samples were cleared in 70% glycerol in 1x PBS overnight.

### Immunohistochemistry

The following primary antibodies were used: Rabbit-anti-GFP A6455 (Life Technologies: used at 1/1000), Mouse-anti-GFP 12A6 (Developmental Studies Hybridoma Bank: used at 1/100), Rabbit-anti-Mesh (Izumi et al., 2012: used at 1/1000), Mouse-anti-Discs Large 4F3 (Developmental Studies Hybridoma Bank: used at 1/50), Mouse anti-Armadillo N27A1 (Developmental Studies Hybridoma Bank: used at 1/50), Mouse-anti-Prospero (used at 1/100), Rabbit-anti-βgal #55976 (MP Biomedicals: used at 1/1000). Secondary antibodies were all highly cross-absorbed varieties: Goat anti-Rabbit Alexa 488 (Jackson Laboratories, used at 1/100), Goat anti-Mouse Alexa 488 (Jackson Laboratories, used at 1/100), Goat anti-Rabbit Alexa 647 (Jackson Laboratories, used at 1/100), Goat anti-Mouse Alexa 647 (Jackson Laboratories, used at 1/100). Stains used included: TMR red (Roche, used according to manufacturer instructions), Phalloidin-iFluor 594 ab176757 (Abcam, 1/1000), and DAPI (1/1000). All confocal images were acquired on a Nikon A1R confocal microscope.

### Analysis of fixed imaging data

#### 1. Ilastik image segmentation

A set of midguts with subjectively standard appearance EB cells (ranging from spindle shape with intense GFP fluorescence to large and round with reduced GFP signal) were used to train a pixel classification project using the ilastik software (version 1.2.2). All object features (Color/Intensity, Edge, Texture) were set to σ=1.6px. 4 training labels were used: Nucleus (blue), EB cytoplasm (green), EB nucleus (red) and Background (grey). The classifiers were trained by painting the sections of the image corresponding to the selected object class. Training involved multiple images, and different z planes and reconstructed cross-sections were used to improve segmentation quality and accuracy. Simple segmentation images were outputted as unsigned 8-bit data in multipage tiff format. All images were processed through batch processing.

#### 2. Morphometric analysis

Two dimensional projections were used for the morphometric analysis. Simple segmentation images were converted into maximum projections in a multichannel RGB format. A region of interest for analysis was assigned using the polygon tool. All files were processed through a custom python macro that outputs EB cells’ morphometric and distribution measurements. Morphometric features were assigned using the “Analyze Particles” tool and items smaller than 5 μm^2^ were excluded from the analysis. The “Measure” function was used to calculate area and shape parameters, such as size, aspect ratio, and circularity (for definitions of these see the ImageJ documentation). Secondary features such as cell coverage and density were derived from primary feature measurements. In the case of EBs in close proximity to each other, due to the lack of cell boundary markers, ilastik pixel classifications were not able to distinguish individual cells. While this did not affect calculation of average EB area, shape descriptors for individual EBs could not be calculated, so the overall merged cytoplasm shape was used. To aid visualisation of single EBs, GFP-positive regions representing EB cytoplasm that contained multiple nuclei were artificially partitioned using the “3D Watershed” algorithm with the nuclei as seeds. ImageJ macro available upon request.

#### 3. Spatial distribution analysis

For EB spatial distribution analyses, F-function and G-function Indices were calculated with the “Spatial Analysis 2D/3D” plugin. The G-function is based on nearest-neighbour distance between objects, while the F-function is based on distances from arbitrary points in space to the nearest object. Both of these functions are based on cumulative frequency distribution (CFD) patterns. The CFD of a given configuration is compared to the average CFD of random configurations with the same number of objects and space. Parameters were set at nb_points=200 and samples=200. ImageJ macros available upon request.

### *Drosophila* adult midgut ex vivo culturing

Shields and Sang M3 Insect Medium (Sigma S3652) was supplemented with 0.5% Penicillin-Streptomycin (Invitrogen 15140-122) and 2% Foetal Bovine Serum (Sigma F4135) to make up the tissue culturing medium. Dissected midguts were transferred from the cavity block onto a 35mm imaging dish (ibidi 81156) using a pair of slightly blunt tweezers held at the hindgut position. A small amount of 1x PBS solution was transferred along with the tissue, allowing the tissue to be encased in a liquid droplet. A tungsten wire was used to orient the tissue so that the anterior midgut was adhered to the bottom of the imaging dish, and the normal loop-like structure of the tissue was preserved. A P20 pipette was used to remove as much 1x PBS as possible, and, as the liquid was removed, the tissue naturally fell onto the treated surface. Once the tissue was exposed, a drop of melted 3% soft melt agarose (see below) was carefully dispensed on top of the sample to encase the tissue and prevent excessive peristalsis during imaging. The working time of molten agarose was 10-15 seconds, during this timeframe if the tissue appeared to have detached from the coverslip bottom, it was readjusted with the tungsten wire. Once set, the sample was topped up with an additional drop of soft melt agarose. The process was repeated until all samples were placed inside the imaging dish (5-7 samples per dish). Following this, additional agarose was dispensed to form bridges connecting each of the tissue islands as well as to the edge of the dish, to prevent the agarose encased midguts floating off from their original positions during transportation. Finally, 1ml of room temperature culturing medium was added. The entire sample preparation process did not exceed 30 minutes.

### Soft melt tissue stabilising agarose

To make 3% soft melt agarose, 0.3g of 2-Hydroxyethyl Agarose (Sigma A9414) was added to 10mL of room temperature Shields and Sang M3 Insect Medium to produce a slurry. The slurry was then heated to 95°C on a heat block with occasional swirling to allow the agarose mixture to be completely dissolved. The molten agarose was dispensed into 500μl aliquots and kept at 4°C. Before use, each aliquot was melted at 80°C and allowed to cool down until it was no longer hot to the touch before dispensing onto midgut tissues.

### Live imaging of *Drosophila* adult midguts

Ex vivo cultured *Drosophila* adult midgut were imaged using a Nikon Andor WD Spinning Disk confocal microscope. Room temperature was kept constant at 21°C during the imaging session. Room temperature dH_2_O was added to the enclosed imaging stage to maintain humidity and prevent sample evaporation during the imaging session. Time-lapse images were acquired using 1024 x 1024 pixel size, at 20x magnification, with stacks collected over 4 hours at 10 minutes intervals. Total depth of stacks was 50μm with a 1μm step-size. One stage position was set for each midgut sample, covering as much of the anterior midgut as possible. The sample was illuminated with the 488nm and 561nm lasers and the GFP and RFP channels were acquired simultaneously with front and back cameras. The exposure time was 300ms. All images were processed using maximum z-projections (ImageJ).

### Analysis of live imaging data

To quantify dynamic behaviour of EB cells, confocal stacks were processed as follows. A region of interest was specified to include in-focus EB cells. Due to the presence of physiological peristaltic movements of the midgut, all time frames were registered to realign cells using the bUnwarpJ plug-in in ImageJ with the deformation mask set to “Fine”. Changes in cytoplasmic shape (cytoplasmic movement) were quantified by subtracting each frame (t) with the previous frame (t-1) in the GFP-channel, taking the absolute mean grey values, then calculating the average intensity within the region of interest over time. To quantify nuclear displacement, the start and end frames of the RFP-channel were processed using the TrackMate plugin, set to radius=5, threshold=20, subpixel particle-localisation and with median filtering. This produced the average Track Displacement for particles over our 4-hr imaging period. To quantify the spatial pattern changes of EB cells throughout the time-lapsed period, all timepoints were analysed for G-and F-function values within the region of interest and the average taken. Analysis and graph generation was done using GraphPad Prism 9. The statistical method used for cytoplasmic movement and nuclei displacement was Student’s t-test, and for spatial pattern analysis was Kolmogorov-Smirnov test. * *p* < 0.05; ** *p* < 0.01; *** *p* < 0.001; **** *p* < 0.0001.

### Counts and analysis of ISC and EB proportions within the anterior midgut

The total number of ISC and EB cells was counted in *GBE^ReDDM^* midguts using the GFP/RFP signals together with DAPI and Armadillo (Arm) antibody staining to identify cell types. EBs were labelled with membrane GFP and nuclear RFP. ISCs were identified by the strong Arm expression on the cell membrane and their basal position within the epithelium. EE cells were distinguished from ISCs by membrane Arm staining that extended all the way to the apical surface. Both ISC, EB and EE cells are diploid and were distinguished from polyploid ECs by DAPI staining. Counts of each cell type and neighbouring cell arrangements were recorded using the ObjectJ plugin which allowed for accurate cell counts across different z-planes. Counts of ISC and EB proportions were exported from ImageJ and analysed using GraphPad Prism9. The statistical method used was the Student’s t-test. * *p* < 0.05; ** *p* < 0.01; *** *p* < 0.001; **** *p* < 0.0001.

### Dextran Sulfate Sodium (DSS)-induced colitis

To induce intestinal colitis with DSS (MP Biomedicals, colitis grade 36,000-50,000), 10-day old flies were starved for 2 hours before transferring to a vial containing 5-6 pieces of 13 x 13mm size Whatman filter paper soaked in 3% DSS in 5% sucrose. The flies were subjected to overnight feeding prior to dissection.

### Induction of regional enterocyte apoptosis with the DEMISE system

The 45D10-Gal4 (Pfeiffer et al., 2011) was used to induce EC apoptosis in the A1 region of the midgut with the DEMISE system. 5-day old females were subjected to 1h heat shock at 37°C to induce reaper expression. Animals were dissected at 0, 24, and 48-hours post heat shock for the examination of EB responses.

### Paraquat-induced midgut damage

To induce oxidative stress with paraquat (methyl viologen dichloride hydrate 98%, Sigma Aldrich 856177), 10-day old flies were transferred onto a vial containing 5-6 pieces of 13 x 13mm size Whatman filter paper soaked in 4mM paraquat in 5% sucrose. The flies were subjected to overnight feeding prior to dissection.

### High throughput screening of MET regenerative cluster regulators

Potential regulators for MET regenerative cluster formation were screened using the ReDDM system in conjunction with candidate gene RNAi. To ensure reproducible regenerative cluster formation and to identify subtle phenotypic variations, experimental flies were kept on the same food for 3 days and subjected to overnight 4mM paraquat-feeding prior to dissection. All images were acquired at 20x magnification and processed through a similar ilastik image segmentation as described above. 6 training labels were used: New EC (red), new EB cytoplasm (green), new EB nucleus (yellow), late EB cytoplasm (cyan), late EB nucleus (magenta) and Background (grey). Simple segmentation images were converted into maximum projections in a multichannel RGB format. Regions of interest for analysis were assigned using the polygon tool. A custom python script was used to identify regions within the midgut that contained features of an EB MET regenerative cluster. A cluster was identified by a new EC being within 50μm of new and/or late EBs. New ECs, new EBs and late EBs (or pre-ECs) were classified if the cells fit our described shape parameters associated with terminal differentiation: new EC nuclei area ≥ 20.0 μm^2^, new EB cytoplasm aspect ratio ≥ 2.0, late EB cytoplasm aspect ratio ≥ 2.0, and late EB nuclei area ≥ 20.0μm^2^. The output of the script includes summary representative images and other regenerative cluster parameters assigned (i.e. cluster count, EB count, clusters coverage and average EB area). For results quantification, the statistical method used was the Welch’s t-test with Bonferroni correction to eliminate instances of false positives associated with multiple t-tests (Bland and Altman, 1995). * *p* < 0.05; ** *p* < 0.01; *** *p* < 0.001; **** *p* < 0.0001. ImageJ macros available upon request.

## Results

### Wildtype Enteroblast characterisation

To perform cell lineage tracking and visualisation of EB cell morphology, we used the ReDDM technique which exploits the differential half-life of fluorophores in combination with the Gal4/UAS system under gal80^ts^ control (Antonello et al., 2015a). EB cells were labelled using the Notch activity reporter Suppressor of hairless (*Su(H)*)-GBE-GAL4 driving a short-lived GFP and a much longer-lived His-RFP that allows one to follow EBs as they differentiate into ECs and EEs (Figure 1A). Then, using RNA interference we altered target gene expression in EB cells. With this system we focussed on the anterior midgut, finding that EB cells in that region had a lower cell density (Figure 1B) compared to those in the posterior midgut (Figure 1C), which allowed for better identification and characterisation of individual cells (Zhai et al., 2017).

**Figure 1.**
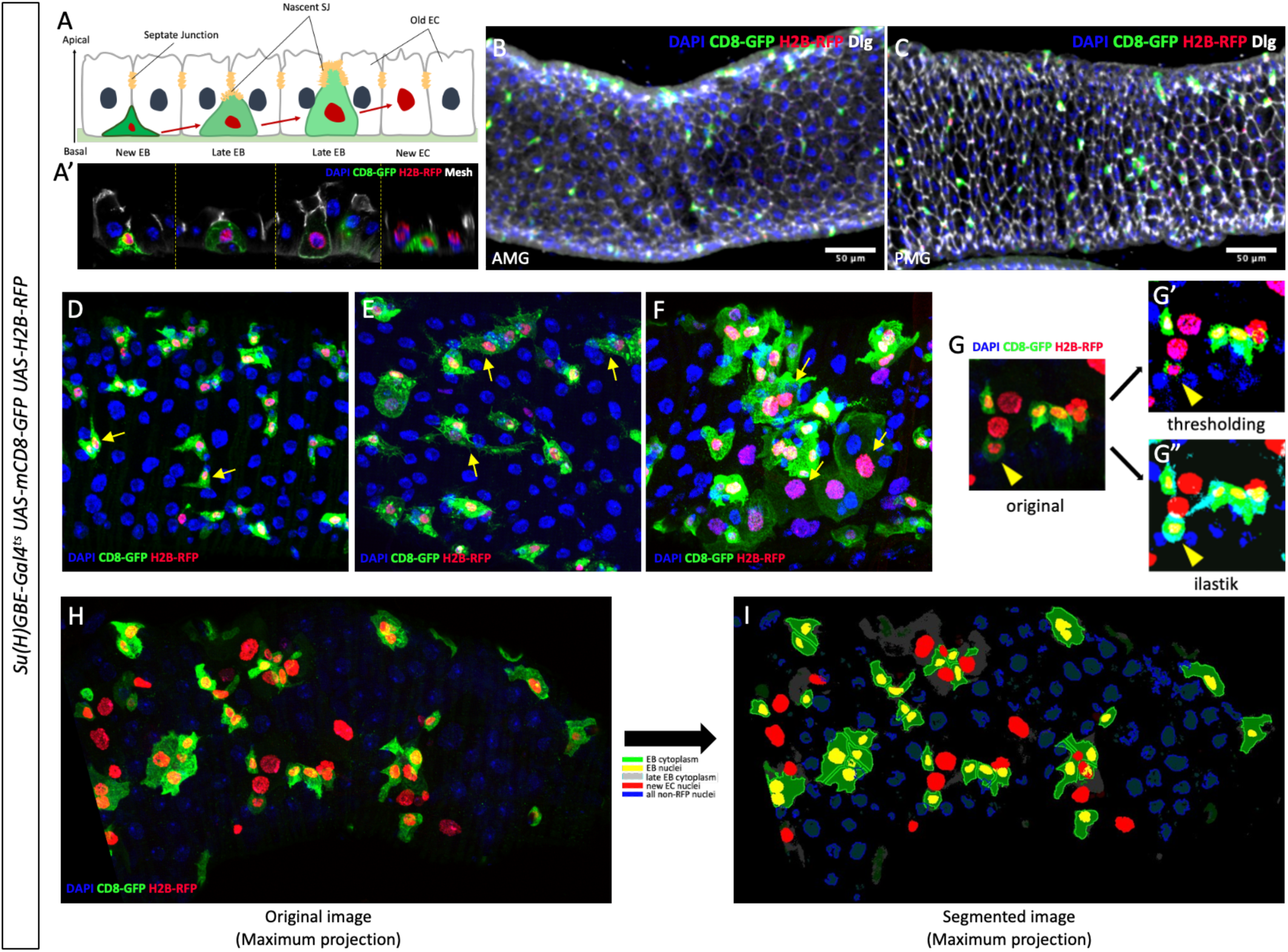
Wildtype EB morphology and image segmentation. (A) EB phenotypic transition during MET visualised using the ReDDM system. Nascent EBs adopt a fibroblast-like morphology and are situated in the basal compartment beneath the SJs of neighbouring epithelial cells (labelled with Mesh; white). When undergoing differentiation and MET, EBs appear to be expanding upwards and have an enlarged cytoplasm, accompanied by an enlarged nucleus, while establishing on their apical surface SJ connections with overlying ECs (marked nascent SJ). Upon completion of differentiation and MET, new ECs form stable SJs with adjacent ECs. (B-C) Anterior midgut (B, AMG) has lower cell density than posterior midgut (C, PMG). Cell junctions labelled with Dlg (white). (D-F) EBs exhibit a wide range of morphologies, as indicated with arrows (N=106): fibroblast-like (D), expanded with extended protrusions (E), significantly enlarged and rounded (F). (G) Image segmentation with the ilastik software results in better background clean up and fluorescent signal identification of cellular components (G”) than simple thresholding (G’). (H-I) Maximum projection of a confocal stack before (H) and after (I) image segmentation.

Initial qualitative assessment of EB morphology through the cytoplasmic GFP staining revealed EBs can take on a wide range of morphologies, ranging from the fibroblast-like morphology typical of mesenchymal cells (Figure 1D), to cells having an expanded cytoplasm with extended protrusions typical of activated EBs (Figure 1E), and finally cells that appeared more terminally differentiated, with a significantly enlarged nuclei, rounded cell perimeter, and showing a reduced GFP signal (Figure 1F).

To relate these cell morphologies with MET, we used antibody staining against key polarity proteins and high-resolution confocal imaging to characterise the incorporation of EBs into the overlying epithelium. It was previously described that quiescent EBs and activated EBs were morphologically different (Rojas Villa et al., 2019). The formation of membrane protrusions is an element of the activation process and begins within a 6-hour window post-tissue damage (Rojas Villa et al., 2019). Here we expand upon the current knowledge and characterise detailed features that distinguish quiescent EBs and activated EBs.

Quiescent EBs were always positioned beneath the Septate Junction (SJ) of neighbouring cells and exhibited typical mesenchymal elongated cellular morphology. In comparison, activated EBs were associated with the cytoplasmic expansion and an upward projection towards the lumen, indicative of cell fate transition as well as active intercalation. We also observed the presence of multiple membrane protrusions in agreement with Rojas Villa et al., (2019), a unique feature to distinguish activated EBs from those further along their differentiation trajectory. At positions of cell-cell contact between an activated EB cell and its neighbours, an enrichment of SJ proteins (i.e. Mesh) could also be observed, which is denoted here as a nascent SJ (Figure 1 A – A’). These observations are in agreement with the Preformed Apical Compartment (PAC) formation stage described in recent publications (Chen and St Johnston, 2022, Galenza et al., 2023).

To fully utilise the morphological information as well as further our understanding of EB-to-EC transition, we next established the morphometric and distribution analysis pipeline. This integrates segmentation of cellular components (using ilastik, a machine learning software), multiple parameters particle analyses (ImageJ), and spatial distribution statistics (Andrey et al., 2010). We found image segmentation using the ilastik software was a superior method of assigning biologically meaningful component identity, compared to simple thresholding based only on pixel intensity (Figure 1G-H) (Berg et al., 2019). The output of the analysis pipeline is a set of parameters that characterise EB morphology and spatial distribution (see Materials and Methods). Key parameters used in the analyses were: circularity, solidity, area, aspect ratio and the F and G spatial distribution indices (Andrey et al., 2010). Additional parameters with biological significance could also be derived (e.g. EB coverage – total EB cytoplasm area/ total area). Using this pipeline, we first established a baseline reference for EB morphology and spatial distribution by analysing cells from wildtype midguts.

Partitioning of cell edges of EBs in close proximity utilised the watershed algorithm (see Materials and Methods). Cellular components are as indicated in key. Late EB cytoplasm (grey) was trained against cells with enlarged nuclei and reduced GFP signal, indicative of reduced Notch signalling down the differentiation path.

### RNAi knockdown of polarity molecules affects EB morphology and MET

To verify that our pipeline could capture phenotypic changes associated with loss of gene function, we selected a few genes with known roles in midgut homeostasis or cellular motility, knocked them down using RNAi, and analysed the effects on EB morphology and spatial distribution. The genes tested were Septate Junction components *Mesh, Snakeskin* (*Ssk*), and *Discs Large* (*dlg*); an integrin subunit, *βv integrin* (*Itgbn*); and a regulator of epithelial integrity, *Sterile 20-like kinase* (*Slik*) (Hipfner et al., 2004; Izumi et al., 2012; Maynard et al., 2010; Okumura et al., 2014; Yanagihashi et al., 2012). Amongst these candidates, RNAi knockdown or mutation of *Mesh*, *Ssk* and *Itgbn* in various midgut cell-types has previously been documented to cause proliferation and differentiation defects and aberrant cell morphologies (Chen et al., 2020; Izumi et al., 2019; Okumura et al., 2014). In the case of *dlg* and *Slik*, though clear evidence of their role in EB polarisation and MET is lacking, we could hypothesise their potential involvement in the process based on established gene functions. Dlg is a well-established regulator of cell polarity and is essential for the formation of pleated septate junctions (Roberts et al., 2012; Woods et al., 1996; Woods and Bryant, 1991). Interestingly, a recent study reported that Dlg did not play a role in the polarisation of adult midgut cells (Chen et al., 2018). Finally, Slik can phosphorylate and activate Moesin, an ERM protein key to connecting cortical actin to the cell membrane, which regulates epithelial cell behaviour (Hipfner et al., 2004).

The features selected for the morphometric analyses were: EB area, EB aspect ratio, EB circularity, EB cytoplasmic coverage, EB count and new EC count. These descriptors together provide a quantitative measurement for EB MET status, as well as the extent of cell proliferation and differentiation.

Knockdown of *mesh* and *Ssk* in EBs resulted in larger cells that were rounder, in agreement with past publications (Figure 2A, B). Interestingly, circularity, which measures smoothness of the cell perimeter, revealed a difference between *mesh* and *Ssk* (Figure 2C). Loss of *mesh* resulted in a decrease in circularity that could be a result of increased membrane protrusions, whereas loss of *Ssk* had no effect on EB circularity. Loss of either *mesh* or *Ssk* had no effect on EB coverage, cell count or newly generated ECs (Figure D-F). Knockdown of *dlg* in EBs produced no effect on cell size but significantly elevated circularity and reduced aspect ratio (Figure 2B, C), both features linked to terminal differentiation, indicating that Dlg may have a role in cell shape or cell identity maintenance. Loss of *dlg* also affected EB coverage, a result of a decrease in EB number (Figure 2 D, E). Together, this result suggests that in contrary to a previous report (Chen et al., 2018), Dlg has a functional role in the adult midgut, at least in EB MET. Unexpectedly, based on previous reports that EB-specific loss of *Itgbn* had no phenotype (i.e. only produced a change with EC-specific knockdown), we found *Itgbn*-RNAi midguts showed an interesting shift in EB shape towards being rounder and having smoother perimeters (Figure 2B, C). EB coverage was also significantly reduced, despite there being no difference in average cell size (Figure 2D). We speculate that this is due to a reduction in EB number (though this parameter was not statistically significant, *p* =0.08) (Figure 2E). Loss of *Slik* in EBs produced the phenotype one would expect with an impaired cytoskeleton – small and rounded cells – but there was no effect on cell number (Figure 2A-E). All four genes examined had no effect on production of new ECs (Figure 2F). These results demonstrate the efficacy of the pipeline in assessing gene function that impacts cell morphology, with the ability to identify subtle differences not evident to simple qualitative assessments.

**Figure 2.**
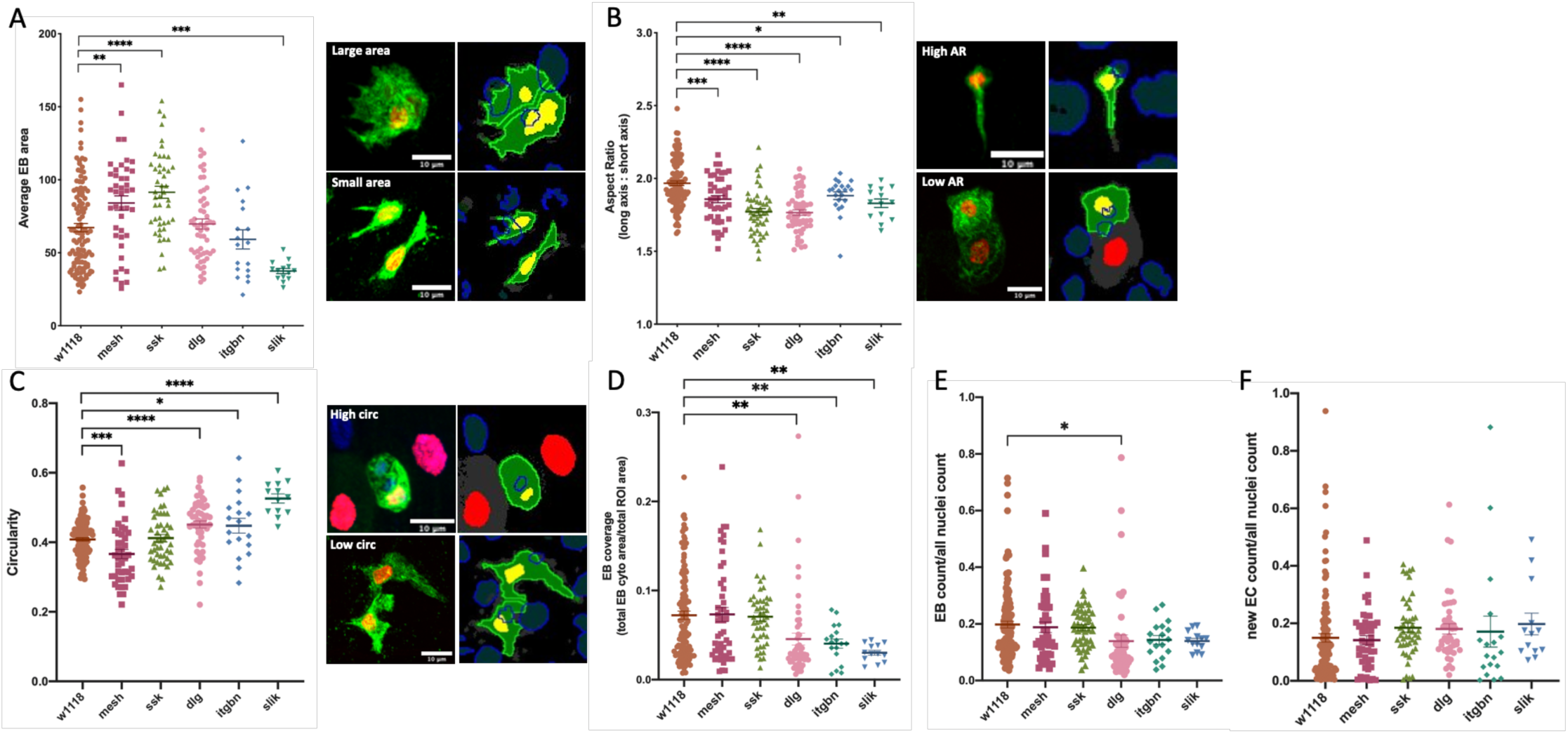
Morphometric analysis reveals differential functional requirements for polarity and adhesion molecules in regulating EB morphology. Targeted gene knockdown in EBs was achieved with *GBE^ReDDM^* crossed to *gene.RNAi* lines and *w^1118^* as control, and hereafter referred to as *gene.IR* or *w^1118^* for simplicity. RNAi knockdown of SJ components *mesh*, *Snakeskin* (*Ssk*) and Discs Large (*dlg*), integrin *Itgbn* and *Sterile 20-like kinase* (*Slik*) in EBs impacted cell size and shape but not the number of new ECs produced. (A) Loss of *mesh* and *Ssk* resulted in larger cells, while the loss of *Slik* produced smaller cells. The cell-cell boundary in this panel was derived using a watershed algorithm to aid visualisation (see Materials and Methods). These artificial boundaries were not used in the morphometric analysis. (B) Knockdown of *mesh*, *Ssk*, *dlg*, *Itgbn* and *Slik* produced cells with a lower Aspect Ratio (AR). (C) Circularity measures perimeter smoothness. The loss of *Ssk* has no impact on smoothness, whereas knockdown of *mesh* led to a rougher perimeter, and loss of *dlg*, *Itgbn* and *Slik* led to a smoother perimeter. As an expanded cytoplasm is a feature of EB differentiation, EB coverage (total EB cytoplasm area/total midgut area assayed) (D) is used as a proxy to measure degree of differentiation, together with EB cell count (E) and new EC count (F) to assess whether our candidate gene knockdowns had an effect on new EB production, level of differentiation, and successful identity transition into new ECs. (D, E) As the proportion of EBs was comparable to that of wildtype when I*tgbn* and *Slik* were knocked down, this excluded the possibility that the significant reduction in EB coverage was due to increased EB number. Hence, loss of *Itgbn* and *Slik* both led to overall EB size shrinkage. Knockdown of *dlg* on the other hand reduced EB coverage as a result of a decrease in EB number. (F) All of the candidates examined when knocked down produced no effect on EBs’ ability to terminally differentiate. N-values: *w^1118^* (123), *mesh* (48), *Ssk* (47), *dlg* (54), *Itgbn* (18), *slik*(13). Blue: DAPI, green: GFP and red: Histone-RFP. Scale bar: 10μm.

### Bimodal distribution pattern is observed for EBs during homeostatic conditions

Having established the importance of assessing morphological features, we wanted to explore the functional significance of the spatial distribution pattern of EB cells. Objects in a confined space can take on different configurations (Figure 3A). We can quantify these different configurations using two spatial distribution functions: the F-function and G-function (Andrey et al., 2010). The G-function utilises nearest-neighbour distance between objects and therefore provides a measure of local positioning. In contrast, the F-function is based on distances from arbitrary points in space to the nearest object and therefore provides a global measure of how evenly distributed objects are within the entire space (see Materials and Methods). To illustrate, one can imagine several scenarios where the spatial pattern deviates from random (Figure 3B): evenly distributed single objects (G=1, F=0), evenly distributed small clusters (G=0, F=0), and uneven distributions of objects (G=1, F=1) or small clusters across the space (G=0, F=1).

**Figure 3.**
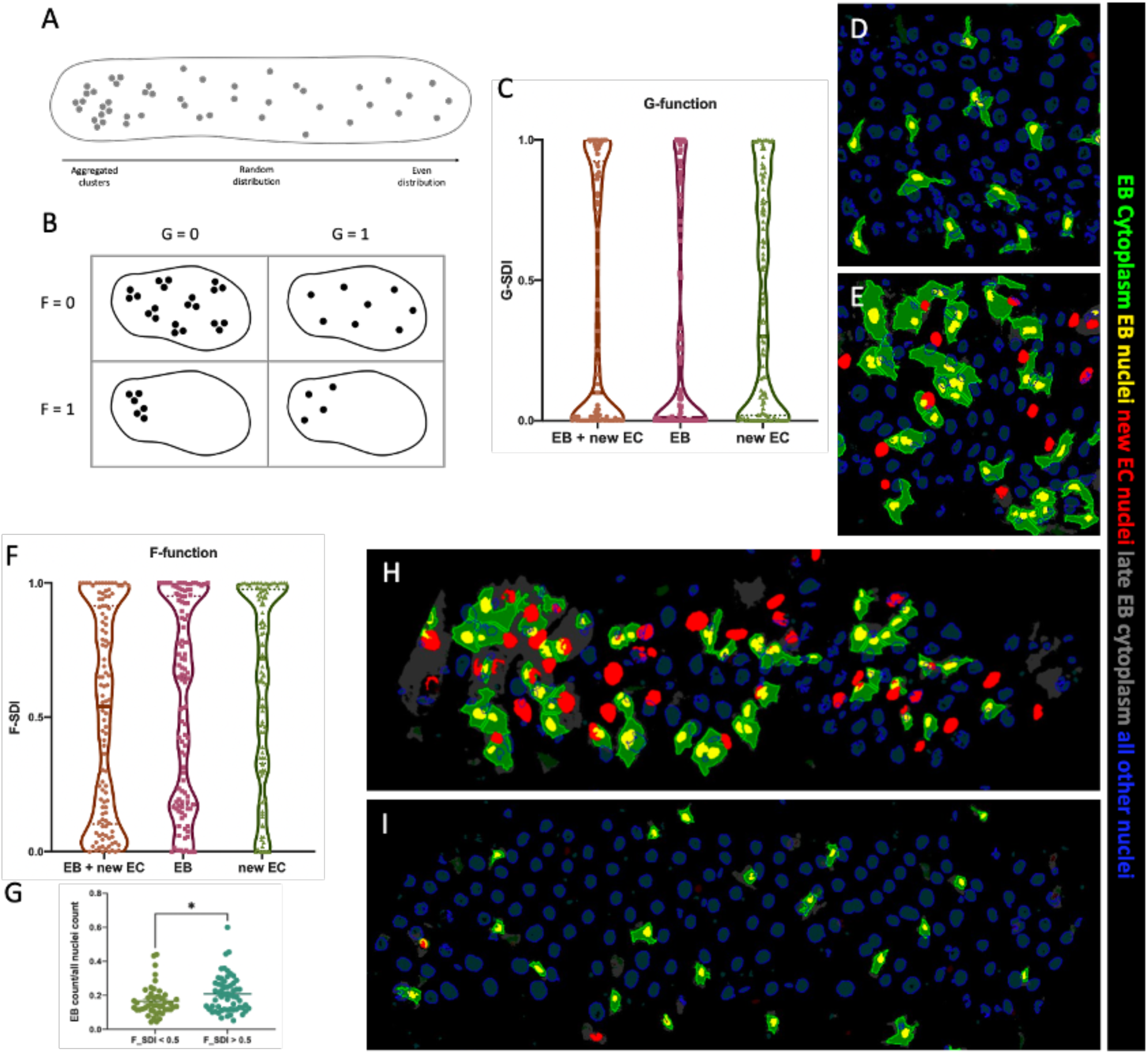
Wildtype EBs exhibit a bimodal spatial distribution pattern in homeostatic conditions. (A) Arrangements of points in space can range from aggregated clusters to being repulsed (i.e. evenly distributed). (B) The combination of two spatial distribution functions F & G can be used to quantify spatial distribution patterns that deviate from a random arrangement. G-function describes nearest-neighbour distance between objects and reflects a local pattern, whereas F-function describes distances from arbitrary points in space to the objects, reflecting a global density variation (see Materials and Methods for details). (C) Wildtype EBs and newly differentiated ECs exhibit a bimodal distribution pattern on a local level (EB: *p* <0.0001; EB + new EC: *p* <0.0001; new EC: *p* <0.0001^#^), reflected with G-function values clustered around 1 or 0. (D) Representative image of a midgut with repulsed EB pattern, G=1. (E) Representative image of a midgut with an aggregated EB pattern, G=0. (F) Wildtype EBs and newly differentiated ECs exhibit a bimodal density variation on a global level (EB: *p* <0.0001; EB + new EC: *p* <0.01; new EC: *p* <0.0001^#^), reflected with F-function values clustered around 1 or 0. (G) F=1 is correlated with an increase in EB cell number, hence total cell count between (H) and (I) differs. (H) Representative image of a midgut with variable EB density, F=1. (I) Representative image of a midgut with even EB distribution, F=0. N=106. Blue: DAPI, green: GFP and red: Histone-RFP. Student’s t-test (* *p* < 0.05). ^#^Comparisons of the G- and F-function distribution patterns with a uniform distribution was performed using the KS test for uniformity.

Although ISCs and EBs have been described as having a random “salt and pepper” or “uniformly scattered” distribution pattern previously (Xu et al., 2011), we uncovered an unexpected bimodal distribution pattern (Figure 3C, F). The distribution graph of the G-function and F-function values of EBs had a dumbbell-like shape, and the top and bottom 10% of the G-function values (G-SDI 2 0.9 and G-SDI σ; 0.1) represented the majority of midguts assayed (15% and 62% respectively). This indicated that most EBs fell under either a “dispersed” pattern or form “aggregated” clusters at a local level (N=106). Midguts with a G-function index of 1 correspond to a repulsed pattern, with isolated EBs (Figure 3D), while those with a value of 0 correspond to an aggregated pattern, with multiple EB cell clusters (Figure 3E). This was also true for new ECs, and EBs and new EC combined (Figure 3C). The bimodal pattern was present on both the local and global scale. At a global level, the F-function describes the pattern of EBs within the midgut epithelium, and therefore reflects cell density. As with the G-function, the same bimodal pattern was observed, though it was less pronounced (Figure 3F). Midguts with an F-function index of 1 exhibit an uneven distribution of EBs across the epithelium, indicative of variable EB density (Figure 3H), while those with a value of 0 have an even distribution (Figure 3I). It is also of note that tissues with a high F-function were associated with an increase in EB number (Figure 3G). Consequently, we were unable to achieve a direct comparison of different density variation in midguts with the same EB count.

### Differentiating EB cells exhibit regenerative cluster pattern of regeneration

Next, we examined if the local pattern of EBs correlated with EB morphology and MET status. When we manually separated out samples into the repulsed and aggregated classes using the G-function cut off point of 0.5 (random pattern), we noted the aggregated EBs coincided with having a global density variation (i.e. high F-index) that was seldom observed in midguts with repulsed EBs (Figure 4A). Midguts with aggregated EBs were also associated with large cluster sizes (Figure 4D). Here we define an EB cluster as a group of EBs in close proximity with membrane contacts (Figure 4H, dotted outline). Since ilastik segmentation is unable to recognise membrane contacts without a plasma membrane marker, it outputs a single cytoplasm containing multiple nuclei. The measurement of cluster size can be extracted by simply counting the number of nuclei within any multi-nucleated cytoplasm. Midguts with repulsed EBs never contained clusters larger than 3 cells in size, with the majority being single EBs or in pairs, whereas midguts with aggregated EBs contained significantly larger clusters in higher proportions (Figure 4D). Cluster sizes varied between individual midguts (Figure 4E, F), but the prevalence of larger EB clusters was always correlated with a reduction in smaller clusters. Interestingly, once we passed the 4-cell mark, larger EB clusters were not any less common than smaller ones (Figure 4G).

**Figure 4.**
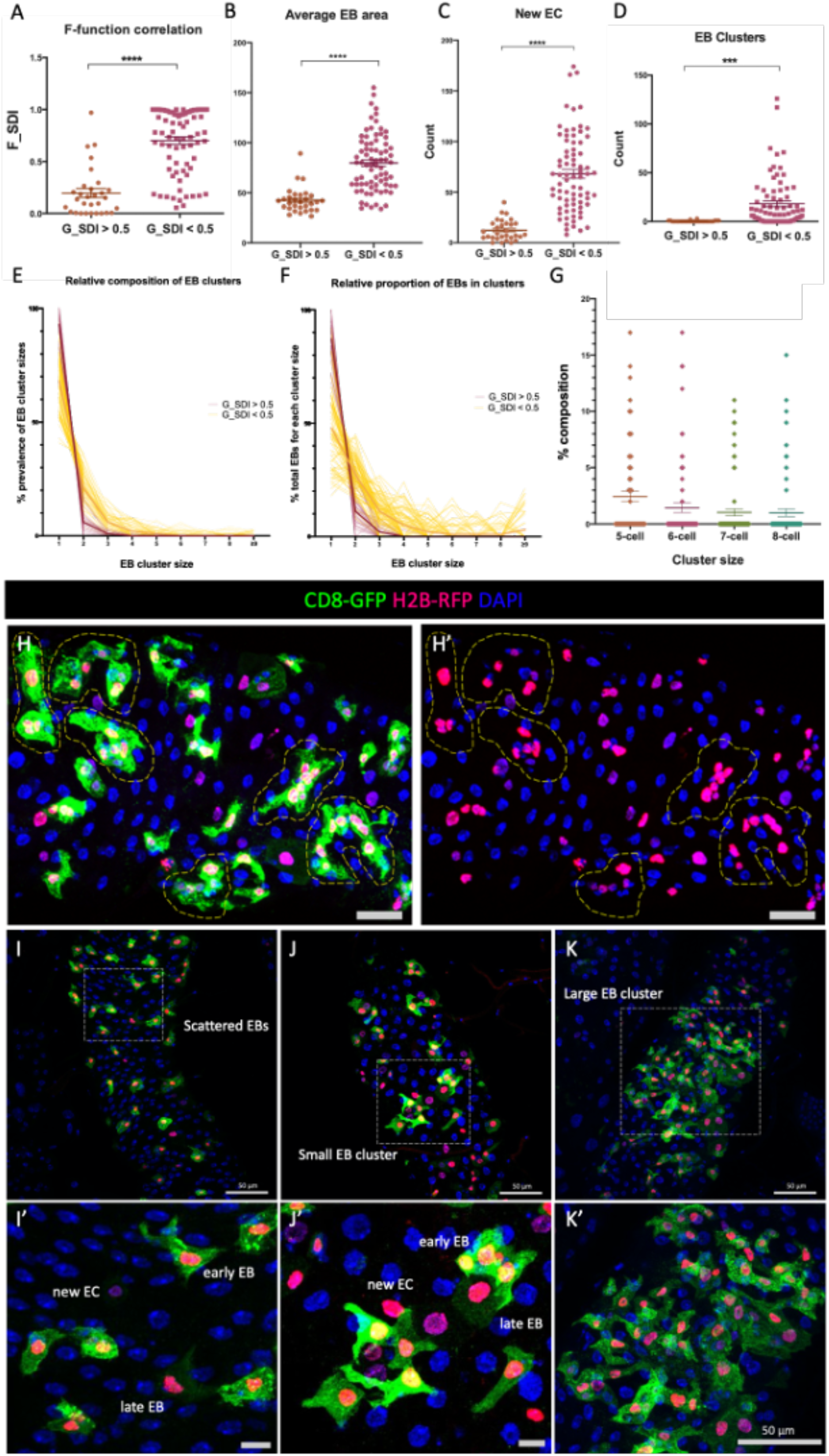
Differentiating EBs exhibit a clustered pattern of regeneration. (A-F) Wildtype midguts with the aggregated EB class (G-SDI < 0.5) vs. the repulsed class (G-SDI > 0.5). N-values: G-SDI < 0.5 (76), G-SDI > 0.5 (30). (A) Midguts with a low G-function value are associated with an elevated F-function value. (B) Average EB size is larger in midguts with aggregated EBs than those with repulsed EBs. (C) Midguts with aggregated EBs contain more new ECs than those with repulsed EBs. (D) EB clusters, classified here as the new EB cytoplasm that contained ≥3 nuclei in ilastik segmented images, were significantly more prevalent in midguts with the aggregated EB class than in the repulsed class. (E) Percentage distribution of EB cluster sizes (i.e. prevalence of single and multi-nucleated cytoplasm). (F) Percentage distribution of EB nuclei that fall within each cluster size category. (G) Data values extracted from (F) from the aggregated EB class, focusing on cluster sizes 5-8. In-depth analysis of EB cluster sizes revealed (E) the majority of EBs exist as singles or in pairs, regardless of their spatial distribution pattern. (E-F) In the midguts with the repulsed EB class, the size of EB clusters never exceeded 4 cells in size (red lines); in the midguts with the aggregated EB class (yellow lines), the prevalence of larger clusters always correlates with a reduction in smaller clusters. Each line represents one midgut, line in bold represents the mean value. (G) Within the aggregated EB class, past the 4-cell size, larger EB clusters are less common than smaller ones. (H-H’) Maximum projection image where various EB cluster sizes were present, large multi-cell clusters highlighted with yellow dotted lines. (I-K’) A spectrum of EB spatial distribution phenotypes: (I) midgut with scattered EBs; (J) midgut with a small EB cluster; and (K) midgut with a large EB cluster. In both (J) and (K), EBs at different stages of their MET process can be observed, hereafter referred as a “MET regenerative cluster”. (I’-K’) Higher magnification images of representative areas highlighted with the white dotted squares in I-K. All data points shown with Mean ± SEM. Mann-Whitney U test (* *p* < 0.05, *** *p* < 0.001, **** *p* < 0.0001).

To test if the aggregated EB pattern was associated with active midgut regeneration, we plotted the number of newly generated ECs with the two local distribution classes. Although almost all midguts contain some new ECs, the midguts with “aggregated” EBs contained significantly more new ECs than those with “repulsed” EBs (Figure 4C). Midguts with “aggregated” EBs were also associated with a significantly higher total EB number compared to that of “repulsed” EBs.

Using the differential half-life of fluorophores and cellular morphology as a proxy for the MET status of individual EBs within a cluster, we can often observe, in a medium size cluster (Figure 4J), the presence of early EBs, late EBs and new ECs, suggesting a local point of regeneration. Such clusters will hereafter be referred to as “MET regenerative clusters”. As MET regenerative clusters contain late EBs, we expect the overall size of EB cytoplasm in regenerating midguts to be larger than those without active regeneration. As expected, “aggregated” EBs had a significantly larger cytoplasmic area compared to “dispersed” EBs (Figure 4B).

### EB cells can exist independently from ISCs

To explore the source of EB “aggregates” formation, we plotted the G-index values of all wildtype midguts sampled against the total EB nuclei counts within. We found that low G-SDIs were associated with a significant increase in total EB number (*p* < 0.05, Student’s t-test). If EB “aggregates” are always associated with increased cell number and differentiation features, we asked whether they were solely a product of local ISC proliferation or could arise via EB cell migration. To that end, we examined if EBs were always associated with ISCs, or if they could exist independently from ISCs. ISCs, EBs, EEs and ECs were identified using a combination of Armadillo staining, which labels ISCs and EBs, *GBE^ReDDM^* driving GFP and His-RFP and DAPI to distinguish diploid vs polyploid cells (see Materials and Methods) (anti-Delta staining, a common marker for ISCs, was unsuccessful in our hands). ISC and EB cells could be characterised into 10 distinct categories of clusters: isolated ISC; isolated EB; ISC/EB pair; ISC/ISC pair; EB/EB pair; ISC/ISC/EB trio; ISC/EB/EB trio; ISC/ISC/EB/EB quadruple; ISC/EB/ISC/EB quadruple and ISC/EB multi-cell clusters of 5 or more cells (Figure 5A-K). Interestingly, 4-cell clusters containing only a single ISC or single EB were not seen. This observation could be explained by the spatial mechanics behind ISC daughter cell fate determination. A 2018 study that analysed real-time ISC division kinetics reported that fate transition is determined by two factors: the horizontal-vertical orientation of the mitotic spindle during division, and whether the parental ISC is flanked by neighbouring EBs (Martin et al., 2018). Simply put, when an ISC is positioned between two EBs, symmetric division is favoured, producing two ISCs and two EBs; instead, when only one neighbouring EB is present, the ISC could adopt any daughter cell fate.

**Figure 5.**
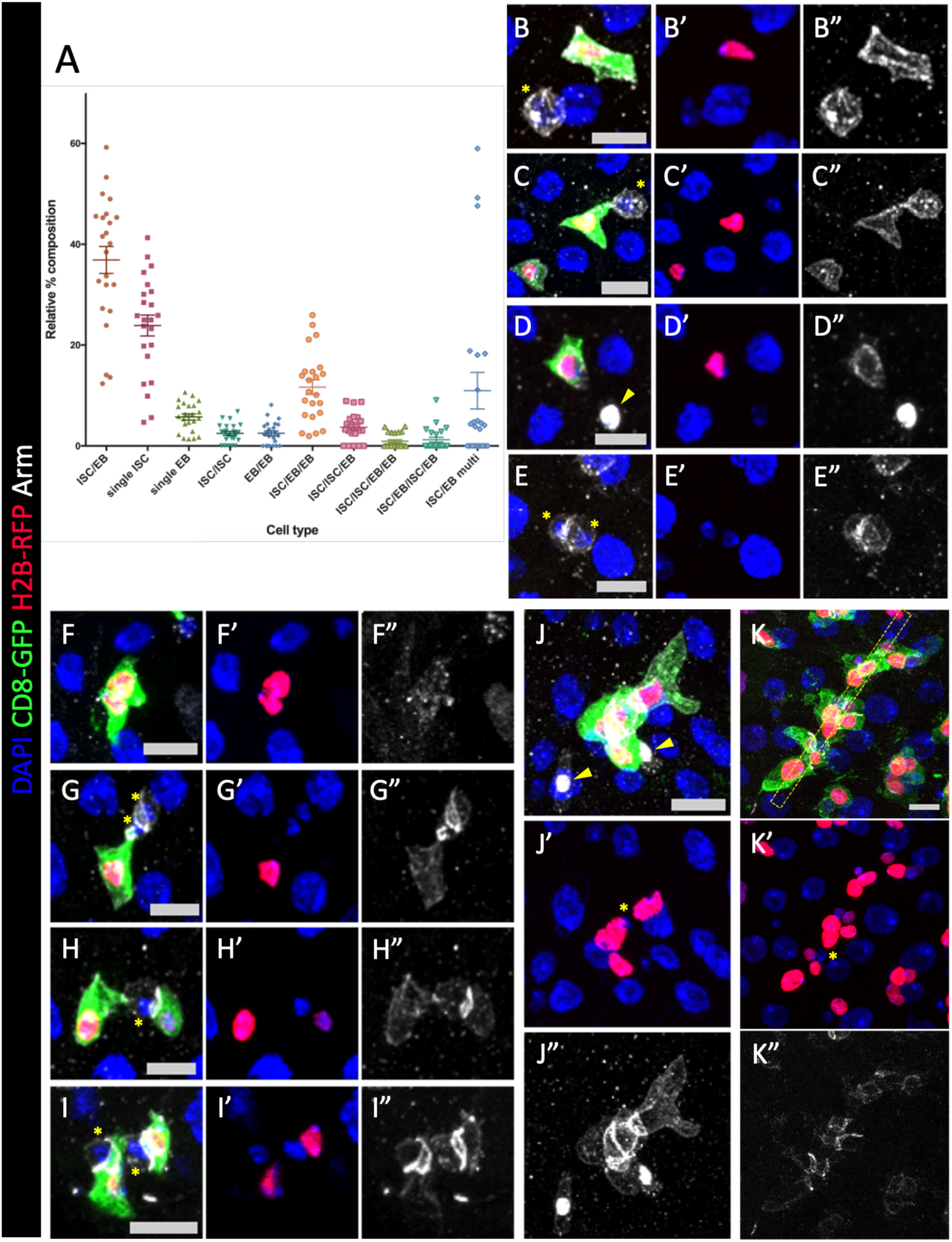
ISC and EB distributions in the anterior midgut range from isolated cells to large cell clusters. (A) Relative percentage composition of cell positioning within a section of the anterior midgut. Each point represents the proportion of ISC + EB that falls into a particular category over the total count of ISC + EB within the assayed area. N=23. (B-K) *GBE^ReDDM^* control midguts counterstained for Armadillo (grey) and nuclear marker DAPI (blue). EB cells are also labelled with membrane CD8-GFP (green) and nuclear H2B-RFP (red). ISCs are identified by stronger Armadillo staining, a small diploid nucleus, and a basal position within the epithelium. Panels showing (B-D) isolated EB with: (B) an isolated ISC, (C) an ISC/EB pair, and (D) an isolated EE; (E) an ISC/ISC pair; (F) an EB/EB pair; (G) ISC/ISC/EB cluster, note enrichment of Arm between ISC and EB cells; (H) ISC/EB/EB cluster; (I) ISC/EB/ISC/EB cluster; and (J-K) ISC/EB multi-cell cluster. ISC: asterisks; EB: RFP-nuclei with GFP-cytoplasm; new EC: RFP-nuclei; EE: arrowheads. All data points shown with Mean ± SEM. Scale bar: 10μm. B-K” are maximum projections.

For multi-cell clusters, the ratio of ISC to EB cells was roughly 1 to 3. The overall ratio of ISCs to EBs within the anterior midgut epithelium was ∼1.3, which contrasts with a previously reported figure of ∼1 for the posterior midgut (Navascués et al., 2012). We also observed that the ratio of single ISCs to single EBs (5.52±0.9) was higher than in the posterior midgut (Navascués et al., 2012). These regional differences may be the result of ISC divisions being less frequent in the anterior midgut (Marianes and Spradling, 2013). The majority of ISC and EB cells existed as ISC/EB pairs (37%) which are likely to be the result of asymmetric divisions of ISCs (Micchelli and Perrimon, 2006; Ohlstein and Spradling, 2006). The next most common category were isolated ISCs (24%) followed by ISC/EB/EB trios (12%). ISC/ISC (2%) and EB/EB pairs (3%), which are likely to arise from symmetrical division of ISCs (Guisoni et al., 2017; Navascués et al., 2012), we found to be much rarer. It is interesting to note that isolated EB cells (6%) were also found (Figure 5B-D), which presumably result from an EB cell migrating away from its parental ISC or the ISC dying after division.

Although the proportions of the various categories of ISC/EB clusters were fairly consistent across different biological repeats, the proportions of multi-cell clusters were highly variable. While most guts had mostly isolated ISC/EBs and small clusters and very few multi-cell clusters, a few guts were clear outliers with several multicell clusters (Figure 5A, J-K). To determine if these midguts might be undergoing regeneration, we correlated the multi-cell clusters with newly generated ECs.

### Wildtype midguts fall into two distinct functional categories

To relate multi-cell clusters with regeneration, we classified the guts into those with new ECs, denoted ‘regenerative’, and those with no new ECs, denoted ‘quiescent’ (Figure 6A-C). As expected, there was a significant difference between the proportions of multi-cell clusters in regenerative vs. quiescent midguts (Figure 6D). In regenerative midguts, we also observed a significant increase in the proportions of EB/EB pairs, while the proportions of single isolated ISCs and ISC/EB pairs were significantly reduced. Hence, we conclude that with regards to progenitor cell distributions, ISC/EBs in quiescent midguts are spaced out and mainly exist in ISC/EB pairs; whereas in ‘regenerative’ midguts, progenitors are present as large ISC/EB multi-cell clusters. We infer that during regeneration, additional EBs are produced from the ISC pool to form the multi-cell clusters in conjunction with coalescence of ‘stock’ EBs. This was supported when we measured the total ISC to EB ratio in ‘quiescent’ (1.53±0.07) vs. ‘regenerative’ midguts (0.75±0.1), where the number of EBs approximately doubled.

**Figure 6.**
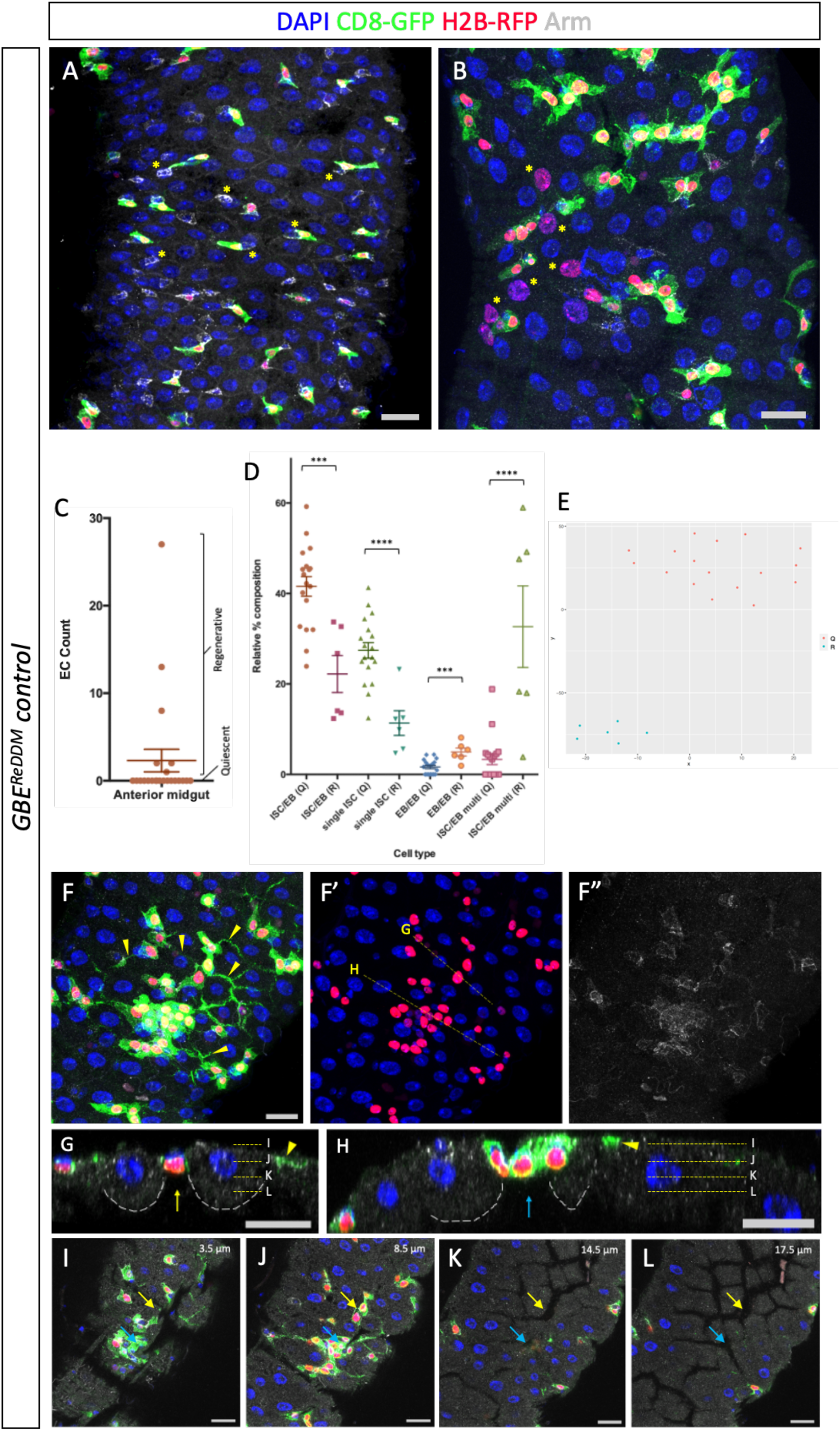
Wildtype midguts fall into two distinct spatial/morphological categories: quiescent and regenerative. (A) A quiescent midgut with spaced out ISCs (asterisks) and EBs (green with red nuclei), often as ISC/EB pairs. New ECs are absent. (B) A regenerative midgut with many newly generated ECs (RFP-positive polyploid nuclei - asterisks). (C) Total new EC count from 23 midguts. Midguts with 0 new EC are categorised as quiescent and those with new EC count ≥1 are categorised as regenerative. (D) Proportions of single isolated ISCs and ISC/EB pairs are significantly reduced, while proportions of EB/EB pairs, and ISC/EB-multi cell clusters are significantly increased in regenerative (R) midguts, compared to quiescent (Q). (E) Quiescent (Q) and regenerative (R) midguts visualised on the t-SNE plot were separated into two isolated clusters. The t-SNE plot was produced with parameters being the proportions of the different ISC/EB cluster categories. (F) A regenerative midgut showing extensive EB protrusions (arrowheads) that extend into the intercellular spaces between nearby ECs. (G-H) Single slice reconstructed cross-sections showing ISC/EB multi-cell clusters coincide with gaps between ECs in the epithelium (arrows), adjacent EC cytoplasms are outlined with white dotted lines. Extended protrusions of nearby EB cells marked with arrowheads. Positions of cross-sections are indicated with yellow dotted lines in F’. (I-L) EC gaps (arrows) can also be visualised across the z-direction, shown in G and H. All midguts shown are counterstained with Armadillo (grey) and DAPI (blue). Scale bar: 20μm. All data points shown with Mean ± SEM. Student’s t-test (*** *p* < 0.001, **** *p* < 0.0001).

Next, we asked if these ‘regenerative’ midguts differed in tissue architecture and cellular arrangements to that of normal homeostatic turnover where old ECs are replaced. Examination of the reconstructed cross-sections along the plane of the EB clusters (Figure 6F) revealed a striking association between the presence of multi-cell clusters and a gap in the EC layer (Figure 6G-L). EBs within the cluster also shown extended membrane protrusions (Figure 6F, arrowheads). Based on this observation, we hypothesised that a break within the midgut epithelium triggers a wound-healing type response of neighbouring EBs. As no ectopic damaging agent had been applied, and based off recent report by Reiff et al., 2019, we speculate that bacterial load accumulation in the food contributed to EC damage and thereafter EB regenerative response.

### Enteroblasts have motility and is the driving force for tissue regeneration

Based on data presented thus far, we propose a model in which local EC damage triggers an activation and directional migration of nearby EBs to coalesce at the site of damage (either through diffusible chemoattractant molecules or mechanical cues). To test the effect of cell damage on EBs, we induced tissue damage using Dextran Sulfate Sodium (DSS) and examined EB behavioural differences between DSS-treated, and control, midguts. DSS feeding induces intestinal inflammation similar to human ulcerative colitis (Amcheslavsky et al., 2009) through disruption of basement membrane organisation, specifically targeting Collagen IV cross-linking in the muscles (Pan and Jin, 2014; Rera et al., 2012). DSS also stimulates ISC proliferation through the Hippo pathway (Amcheslavsky et al., 2009). Experimental animals were subjected to oral feeding of yeast paste supplemented with 3% DSS 20 hours prior to processing.

In order to assay EB behaviour, we turned to live imaging. Our understanding of EB motility is rudimentary with conflicting evidence present in current literature (Antonello et al., 2015a; Hu et al., 2021; Rojas Villa et al., 2019). Using an ex vivo culturing set up, we successfully achieved tissue survival for 30 hours post-dissection (Figure 7 A-B). Functional assessment of tissue health was performed through visual confirmation of the presence of midgut peristalsis and crop contractions pre- and post-imaging (Figure 7C, Movie 1). In control midguts, we found that EB cells had limited motility over a 4-hours imaging period (Figure 7D top panel, Movie 2). Both small scale active extension and retraction of membrane protrusions were observed. In DSS-treated midguts, we observed a significant increase in EB cytoplasmic movement (Figure 7E, Movie 3), and formation of 1-2 EB clusters in all guts (Figure 7D, Movie 3). Together, our observations are suggestive of directional movement to form regenerative clusters. Despite the increase in cytoplasmic movements, we did not observe any variation in nuclei displacement (Figure 7F).

**Figure 7.**
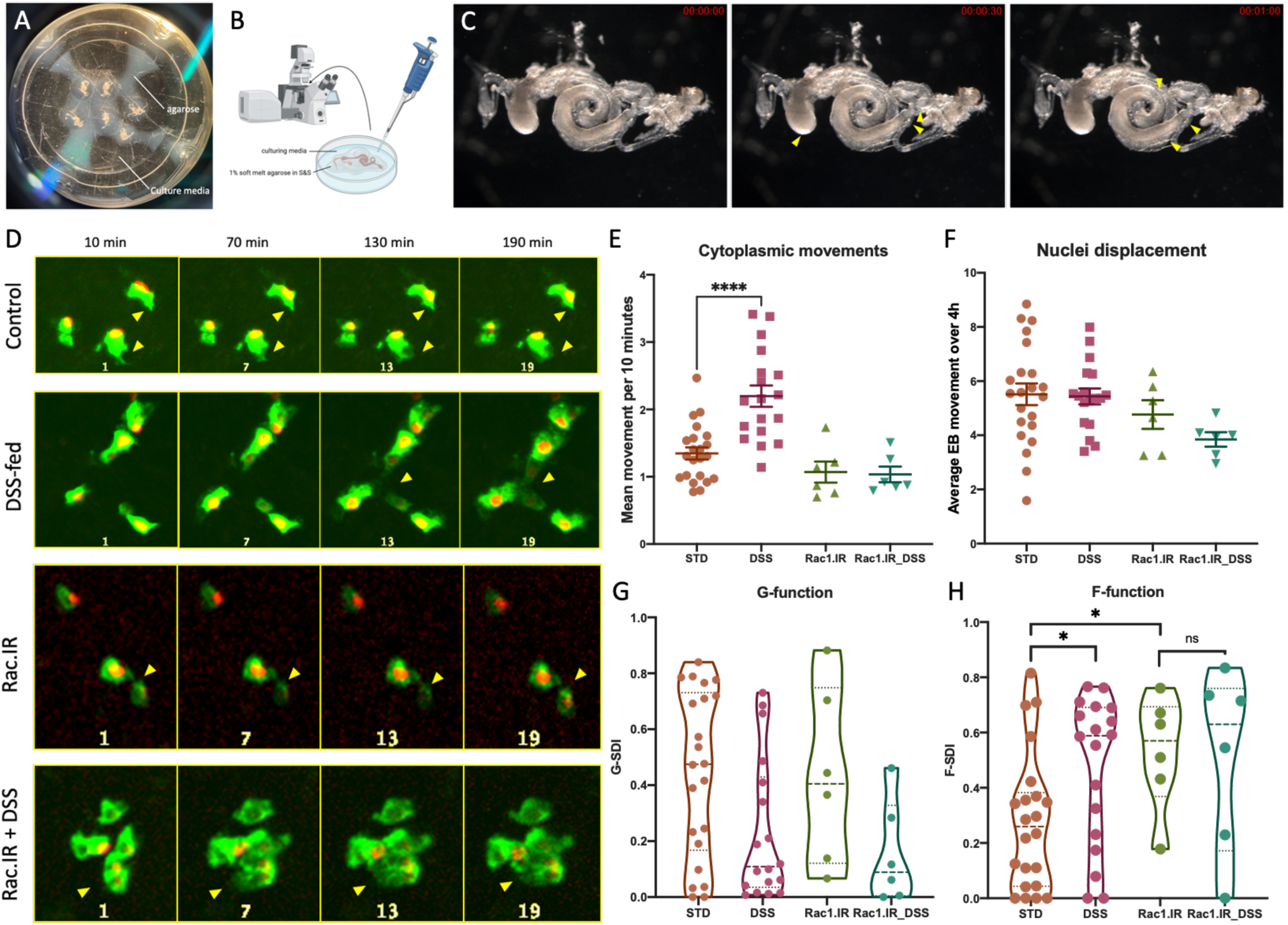
Live imaging of EB cells in the adult midgut. (A-B) ex vivo imaging set up. Freshly dissected midguts were encased in soft melt agarose and cultured in modified Shields and Sang insect media in an ibidi dish. Additional culture media was supplied just prior to imaging with a spinning disk microscope. (C) Midgut tissue visualised with Leica M165 microscope post imaging session. A 1-minute duration video was recorded and peristaltic movements of the midgut, as well as crop contractions were still observable (arrowheads). (D) Differential membrane activities of EB cells subjected to control, DSS-treatment, RNAi knockdown of *Rac1*, and *Rac1*-deficient EB cells with DSS-treatment (arrowheads). EB cells subjected to DSS-treatment appeared to elongate and form membrane contacts. *Rac1.IR* EBs appeared more rounded with limited membrane movements. DSS-induced cytoplasmic elongation was limited in *Rac1.IR* EBs. (E-F) Quantification of mean cytoplasmic movement and nuclei displacement across all EB cells within the region of interest (see Materials and Methods). Each point represents one midgut sample. All data points shown with Mean ± SEM. Student’s t-test (**** *p* < 0.0001). (E) DSS-feeding significantly increased membrane motility in control EBs, and such behaviour was abolished by the knockdown of *Rac1*. No significant difference was observed between the wildtype control and *Rac1* knockdown controls. Unit: Mean Grey Value. (F) No differences in nuclei displacements were observed in the wildtype group as well as the *Rac1* knockdown group over the 4-hours imaging period. Unit: Pixels/10 min. (G-H) Mean EB spatial distribution indexes in control, DSS-treatment, RNAi knockdown of *Rac1*, and *Rac1*-deficient EB cells with DSS-treatment groups. All data points shown with violin plots. Kolmogorov-Smirnov test (* *p* < 0.05). (G) Mean G-function values representative of local EB distributions suggested the presence of more EB aggregates in both the DSS-treated wildtype as well as *Rac1* knockdown groups compared to the controls, though not statistically significant. (H) Mean F-function values representative of the overall EB distribution pattern within the midgut has shown a significant density variation between wildtype EBs with those in midguts treated with DSS. Global density variations can also be observed when *Rac1* is knocked down compared to wildtype. The number of samples assayed: wildtype with control diet (N=22), wildtype with DSS (N=18), *Rac1.IR* with control diet (N=6) and *Rac1.IR* with DSS (N=6).

If EB migration is necessary for cluster formation, one could imagine cells with defective motility would be unable to form clusters. To disrupt the actin cytoskeleton of EB cells we used RNAi knockdown of the small GTPase *Rac1*. We found that the baseline cytoplasmic movement was slightly reduced (Movie 4), but more significantly the increased cytoplasmic movement response to DSS was abolished (Figure 7E, Movie 5). However, small EB clusters were still occasionally present in the samples analysed (e.g. Figure 7D, bottom row) from the first time point of imaging. Since we did not observe active formation of clusters, we propose these clusters had formed prior to imaging.

To complement our qualitative assessment that EBs can form clusters and become more aggregated with DSS, we performed spatial distribution analysis on the time-lapse data. A single spatial distribution index (G-SDI and F-SDI) was produced at every timepoint of acquisition. Using the mean values of all 24 timepoints in each sample, we found no statistical variation of the G-function for EBs from all four treatment groups (Figure 7G), but found significant variation in the F-function between DSS-treatment and RNAi knockdown of *Rac1* when compared to controls (Figure 7H). Together, this suggests that the EB global density variation is partly due to cell motility but may also include local ISC asymmetric proliferation.

In addition to damage induced by DSS, we adapted an existing genetic method to create mosaic intestinal damage within a confined region of the midgut and evaluated EB responses (Figure 8D). Using the DEMISE system (Dual Expression Method for Induced Site-specific Eradication, Cohen et al., 2018), we can obtain temporally and spatially defined EC apoptosis in the anterior midgut with inducible *UAS-reaper* (*rpr*) expression under the regulation of *GMR45D10-Gal4* (hereafter *45D10* for control, and *45D10>DEMISE* for experimental). The expression profile of GMR45D10-Gal4 is in agreement with past publication (Figure 8A, A’) (Marianes and Spradling, 2013). EBs were labelled with *Su(H)GBE-LacZ* expression.

**Figure 8.**
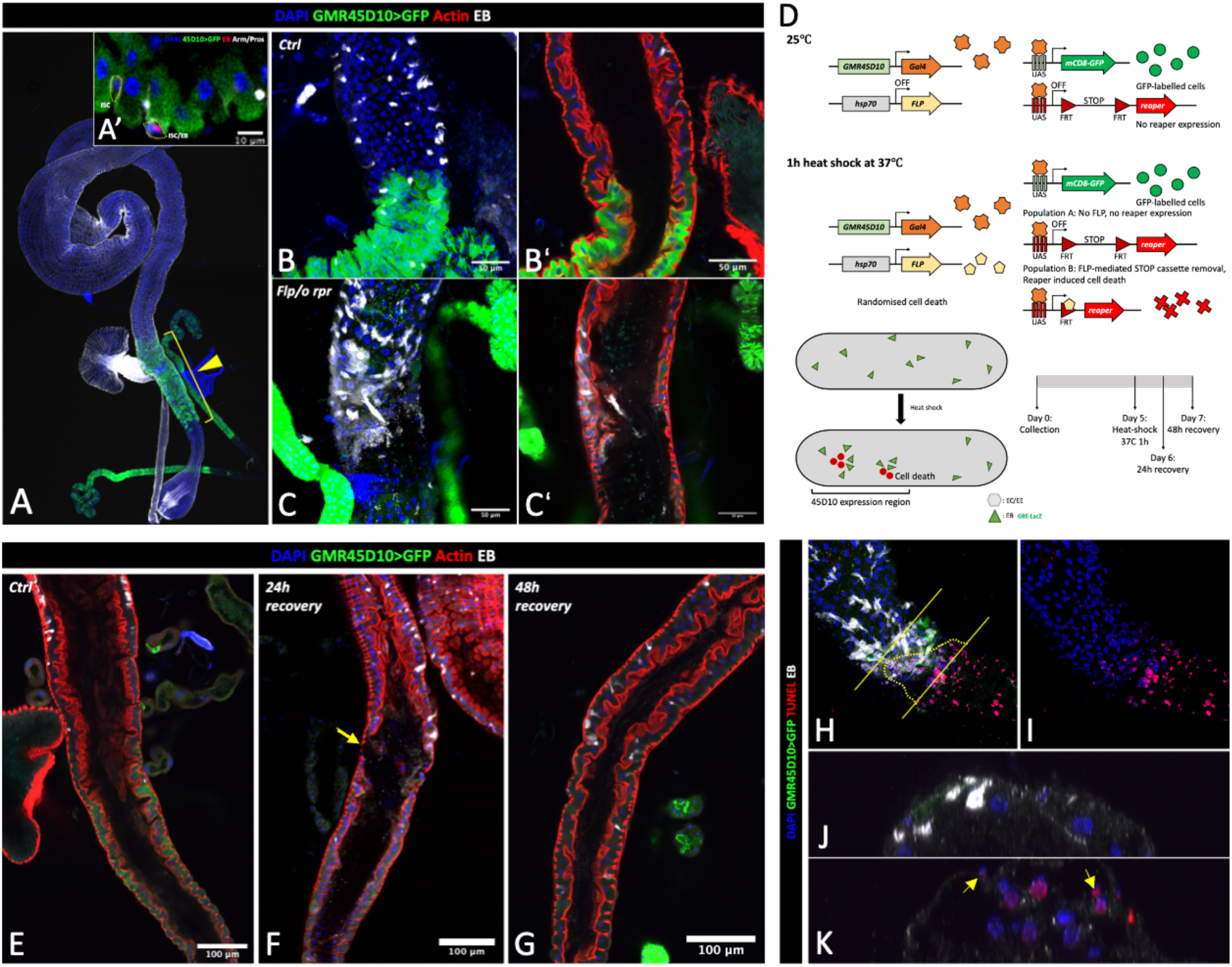
Genetically induced large-scale EC loss could be restored within 48 hours. (A-A’) The expression domain of the GMR45D10 enhancer (arrowhead) is located at the A1 region of the adult midgut and does not target diploid progenitor cells. (B-C) Cell death induced in the 45D10 region using the DEMISE system produces a significant disruption in epithelial structure and changes in EB distribution and morphology. (B) Control midgut 24 hours post-heat shock. N=29. (C) *45D10>DEMISE* midgut 24 hours post-heat shock. N=41. (D) Schematic representation of cell death induction with the DEMISE system in the 45D10 region and the expected outcome. Using a clever combination of the Gal4/UAS and the FLP/FRT system, the DEMISE system was devised to allow for an induction of randomised cell death in conjunction with regional target gene manipulations through temperature regulation. In our setting, the only UAS transgene used was *UAS-GFP* for 45D10 domain labelling, which occurs at 25°C. Mosaic cell death within the region was induced with 37°C heat shock, which led to a portion of cells synthesising the Flippase protein with subsequent removal of the *FRT-Stop-FRT* cassette, allowing for reaper expression. From our working model, if EB directional migration occurs in response to damage, we expect to find a redistribution of EBs in that they are clustered around the domain of damage and exhibit reduced prevalence in the adjacent zone. Flies subjected to heat shock induction were allowed to recover for various amounts of time (in this case showing 24 and 48 hours) prior to examination. (E) A representative control midgut at 24 hours post-heat shock with regularly shaped epithelial cells and a continuous epithelium. (H) Significant disruptions to the midgut epithelium in *45D10>DEMISE* flies 24 hours post-heat shock. Gaps within the epithelium are present (arrow) and cell remnants are present within the lumen. By 48 hours (G), the cellular arrangement of the epithelium appears indistinguishable from controls (E). N=17. (H-K) Tissue damage produced by the DEMISE system occurs through cellular apoptosis. Apoptotic cells are marked with TUNEL staining and appear to be restricted in the 45D10 domain (J versus K). TUNEL-positive cells are found inside the midgut lumen (K). N=12. Panel B-C and E-G are true cross sections, J-K are reconstructed cross sections. Scale bar as presented within the figure.

Initial assessment of *45D10>DEMISE* midguts allowed to recover for 24 hours showed the GFP signal of Gal4-expressing ECs was greatly reduced compared to the control (Figure 8B, C), a likely outcome from the removal of previously GFP-positive cells. As this made clear visualisation of damage boundary difficult, we used the presence of salivary glands adjacent to the A1 compartmental boundary as an additional independent reference point to locate the damage site. At 24 hours post-heat shock, in the absence of damage, EBs exhibited a dispersed pattern (Figure 8B, L). Consistent with our earlier results, when *rpr* expression was induced, we observed an increase in EB size and tissue coverage, as well as the presence of EB cluster (Figure 8C, M). Phalloidin staining, which labelled visceral muscles (VM) and microvilli, revealed in clear contrast to control, a significant disruption in epithelial integrity whereby the EC-layer in the 45D10 region appeared to be largely lost, presumably due to excessive EC shedding (Figure 8B’, C’). The VM staining did not appear visually different, suggesting the underlying musculature was unaffected by EC loss.

Next, we confirmed that EC loss in the *45D10>DEMISE* midguts was indeed due to apoptosis using TUNEL staining, which labels double-stranded DNA breaks that occur at a late stage of apoptosis (Figure 8H-I, K). As expected, apoptotic ECs were restricted to our defined region. Interestingly, some TUNEL-positive smaller nuclei were also present, suggesting the other cell types were also undergoing apoptosis (Figure 8K, arrow). Furthermore, the EB cluster was found near to the boundary of the TUNEL-positive cells, and EBs further away from the damaged region exhibited an elongated morphology (Figure 8H, dotted region, arrows). This result further supports the hypothesis that directed EB migration is associated with post-damage midgut repair.

Finally, when *45D10>DEMISE* midguts were allowed to recover for 48 hours, the continuous epithelial architecture appeared completely restored (Figure 8G), as the break in the epithelium seen at 24 hours (Figure 8F, arrow) was no longer present. All animals remained alive throughout the duration of the experiment (data not shown), indicating large-scale EC apoptosis does not lead to lethality.

### Testing gene candidates for MET regenerative cluster formation and directional motility

To better understand cluster formation in tissue repair it would now be of interest to conduct a large-scale screen for genes that may be involved. To facilitate this, we developed a more rapid screening approach and trialled it with a small candidate-screen of cell surface receptors and mechanosensors. For the screen, we optimised the sample preparation method to forego high resolution imaging with immunostaining, and instead used only the endogenous fluorescence signal of the *GBE^ReDDM^* flies with improved tissue clarity through 5% sucrose solution feeding overnight. Although this approach reduced our ability to visualise fine morphological details (i.e. thin membrane protrusions), it reduced sample preparation time from 3 days down to 1 day, thereby making it suitable for efficient screening.

Next, we established baseline conditions in which the EB clusters were consistently being generated at a level of approximately 2 clusters per midgut, which would allow detection of suppressors and enhancers. Two independent methods for achieving increased MET regenerative clusters were utilised: keeping the flies on the same food for an extended period of time (Reiff et al., 2019), which is thought to allow build-up of microbes and subsequent EC damage (Apidianakis and Rahme, 2011) and oral supplementation of paraquat. We found we could consistently generate 1 or 2 medium sized MET regenerative clusters with 4mM paraquat in conjunction with food change every 3 days.

Next, a custom python script was written to objectively classify the regenerative clusters (see Materials and Methods) defined as being a cluster of EB cells in the vicinity of new ECs. The identification of MET regenerative clusters takes into account the presence or absence of new ECs, as well as the proximity and morphological features of EBs surrounding them (Figure 9A).

**Figure 9.**
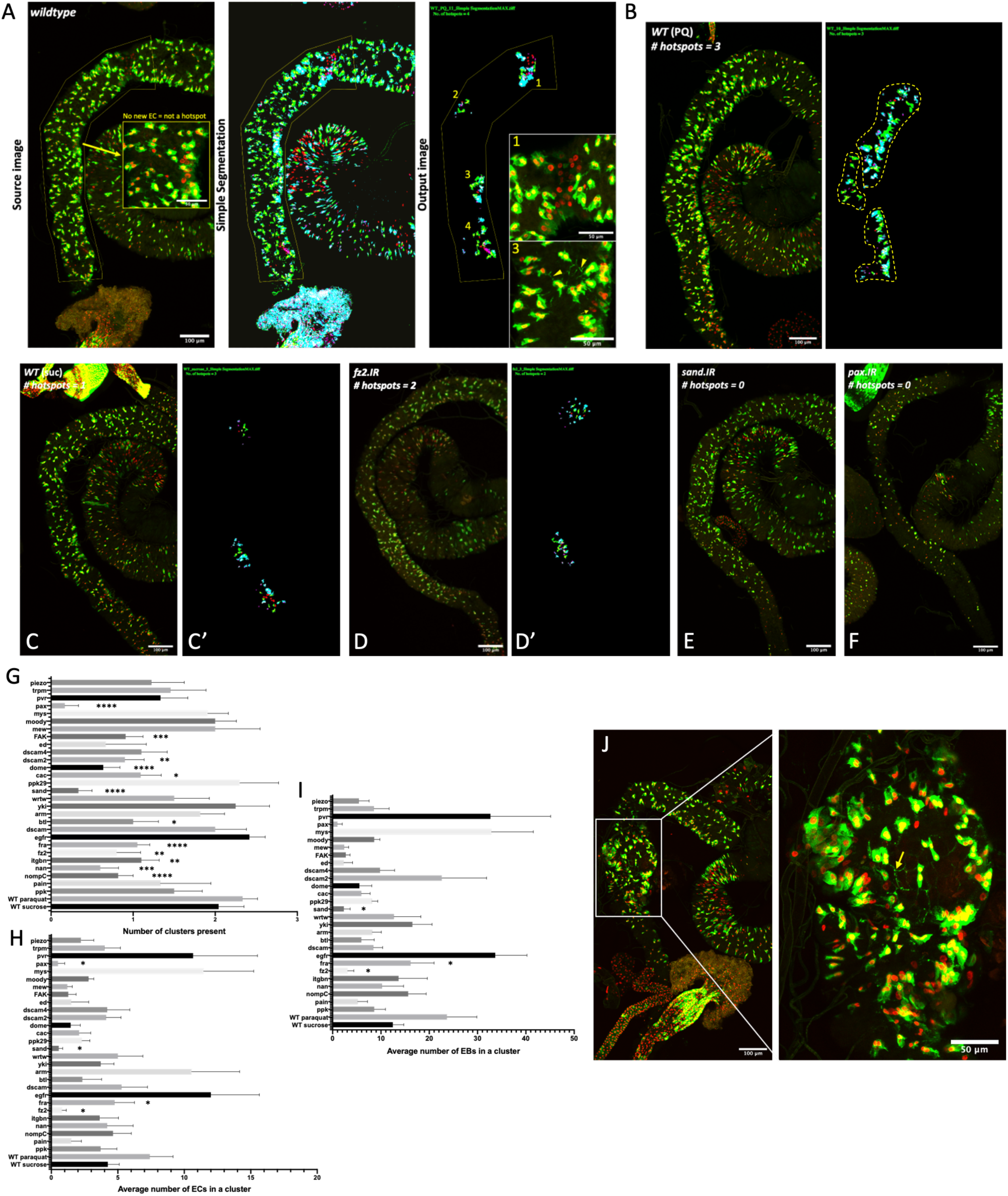
High-throughput screening of potential regulators of MET regenerative cluster formation. (A) Custom python script workflow for the identification of regenerative clusters (see Materials and Methods). (B) Representative image of wildtype midguts with PQ treatment. Clusters identified with the python script are shown in (B’). (C) Representative image of wildtype midguts with sucrose treatment. Clusters identified with the python script are shown in (C’). (D) Representative image of midguts containing *fz2* knockdown EBs with PQ treatment. Clusters identified with the python script are shown in (D’). (E) Representative image of midguts containing *sand* knockdown EBs with PQ treatment. No clusters identified. (F) Representative image of midguts containing *pax* knockdown EBs with PQ treatment. No clusters identified. (G-I) Quantification of cluster parameters used in the analyses of the candidate gene knockdown effects: (G) The number of regenerative clusters present; (H) Average number of new ECs within a regenerative cluster; and (I) Average number of new EBs within a regenerative cluster. (J) A wildtype midgut with an incidental bulge associated with EBs with extensive protrusions (arrow) and local MET regenerative clusters around the periphery. All images shown as maximum projections. All data points shown with Mean ± SEM. Welch’s t-test with Bonferroni corrections (* *p* < 0.05, ** *p* < 0.01, *** *p* < 0.001, **** *p* < 0.0001). Scale bar: 100μm.

As a proof of concept, we selected a small list of putative mediators of MET regenerative cluster formation and applied them to our screen through EB-specific RNAi knockdowns. We hypothesised that the formation of clusters is mediated by mechanotransduction and/or diffusible ligands. Therefore, we selected some mechanosensors based on their differential mRNA expression in EB cells vs. other cell types using data from FlyCellAtlas (Li et al., 2022) and a number of cell surface receptors based on known biological functions (Table 1). Each of the gene knockdowns were compared to the paraquat group.

**Table 1.**
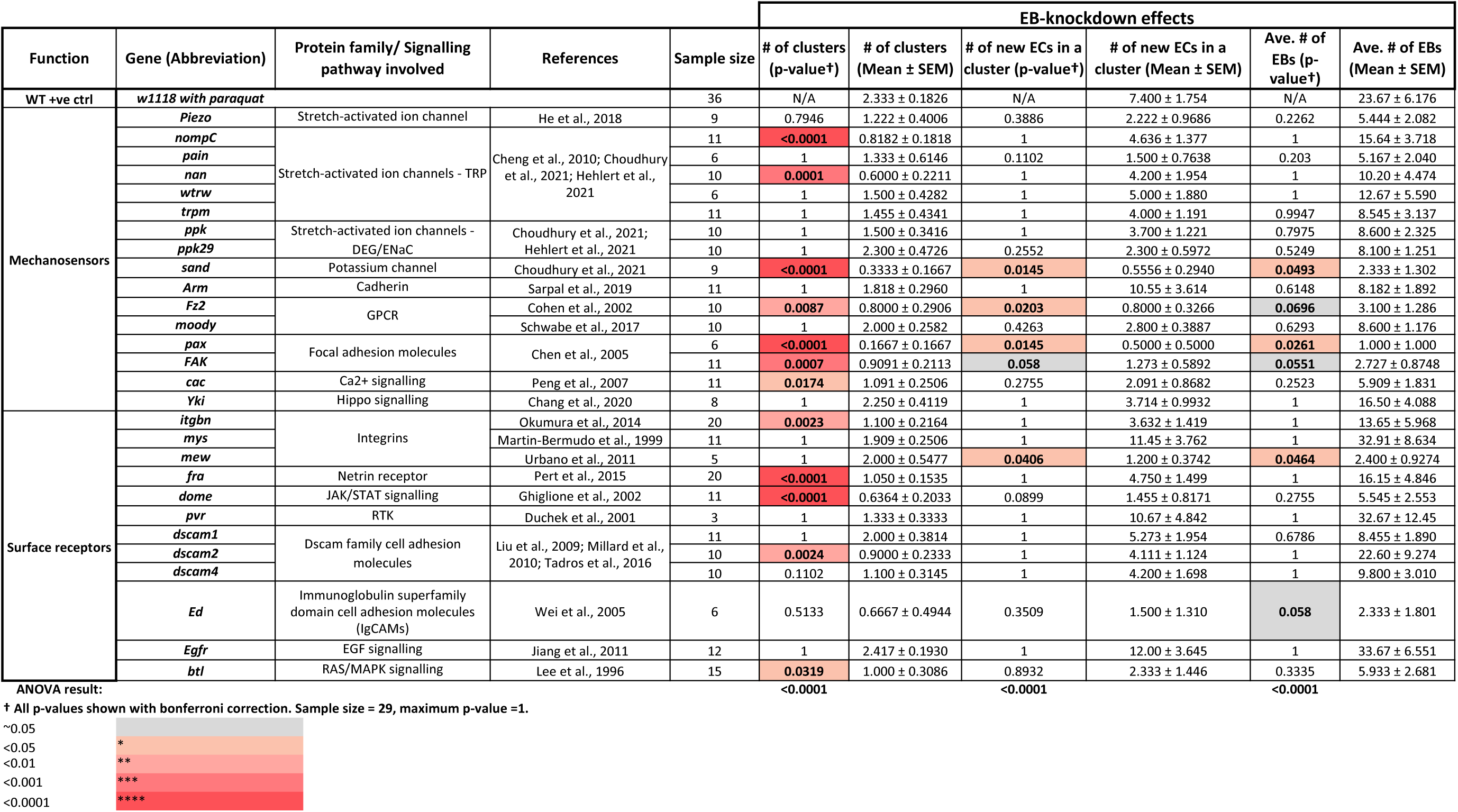
Regenerative cluster analysis of potential regulators. All p-values presented were calculated using the Welch’s t-test with Bonferroni correction applied. Significant p-values are colour coded as per legend. Sample size for each of the gene candidates are as indicated within the table.

We first looked for candidates that significantly altered both the number of clusters generated as well as the number of EB and ECs found within the clusters. Three candidates fit within this category: two-pore domain K^+^ (K2P) channel subunit *sandman* (*sand*), G-protein-coupled receptors *frizzled 2* (*fz2*), and focal adhesion adaptor protein *paxillin* (*pax*). Using the number of new ECs identified as a measurement for regeneration, *sand*, *fz2*, and *pax* are likely to be required for both cluster formation as well as regeneration (Table 1, Figure 9D-F, G-I). Next, we isolated candidates with only cluster count variation but no impact on the number of ECs and EBs. This group contained 4 mechanosensors: transient Receptor Potential (TRP) channel family members *no mechanoreceptor potential C* (*nompC*) and *nanchung* (*nan*), Focal adhesion kinase (*Fak*) and a voltage-gated Ca2^+^ channel subunit *cacophony* (*cac*); and 5 surface receptors: integrin subunit *βν integrin*, Netrin receptor *frazzled* (*fra*), JAK/STAT signalling receptor *domeless* (*dome*), Dscam family cell adhesion molecule *dscam2* and EGF signalling ligand *breathless* (*btl*). Out of the 9 candidates, loss of *Fak* in EBs also produced a strong reduction in the number of EBs and ECs, however this result was not statistically significant (Table 1, Figure 9G-I). We can infer from the lack of variation in EB and EC count that the other 8 candidates may only be required for cluster formation and not regeneration. Lastly, loss of integrin subunit *multiple edematous wings* (*mew*) in EBs did not impact the number of clusters within each midgut but led to a significant reduction in the number of new ECs and EBs within a cluster (Table 1, Figure 9G-I). This result is unsurprising as the integrin complex is an essential component to EB-to-EC differentiation and polarisation (Chen et al., 2018, Chen and St Johnston, 2022). Together, we have uncovered several promising candidates that warrant further studies into the mechanosensing and chemotatic properties of EB cells.

Consistent with our idea that EBs can detect mechanical signals, we occasionally observed (in 4 separate instances) midguts with a bulge (perhaps due to stochastic tissue damage) which coincided with an enrichment of EB clusters. EB cells within the bulged region also had elaborate protrusions (Figure 9J, arrow), a phenotype associated with EB activation (Rojas Villa et al., 2019). Regardless of the cause of the bulging, the association of EB clusters and new EC formation suggest that EB may be responding to some form of pressure-induced cell shape changes in the environment.

## Discussion

Using the pipeline, our results revealed a previously unreported bimodal spatial distribution of EBs in the wildtype adult midgut. EBs tended to either have a dispersed local pattern (high G) with even global density (low F) OR an aggregated pattern (low G) with varied global density (high F). These patterns of EB distribution correlated with the formation of newly differentiated ECs: midguts with dispersed patterns had few new ECs whereas midguts with aggregated clusters had more. In addition, the clusters of EBs also exhibited many late EBs and new ECs, leading us to term these regions “MET regenerative clusters”, and to term the two categories of midguts quiescent and regenerative.

Consistent with Antonello et al. (2015), we found that new ECs often exist in clusters. This observation was briefly noted in Tamamouna et al., (2020). The non-homogenous new EC production suggests that regeneration may be responding to a localised signal of demand. Some evidence exists to support this hypothesis, as some signalling pathways necessary for stress responses were found to have a non-homogenous reporter expression pattern along the midgut length when the animal was subjected to infections or stress conditions. These include 10XSTAT-GFP and upd-lacZ (Zhou et al., 2013) and dpERK (Houtz et al., 2017). There is also evidence of cell death clustering based on a caspase sensor (Arthurton et al., 2020).

An important question is, how do MET regenerative clusters arise? Are they due to the active migration of EBs to the point of damage or a product of local ISC divisions and differentiation? There is some evidence that EBs have motility (Antonello et al., 2015a; Rojas Villa et al., 2019), but it is unclear if they are able to move away from their parent ISC. To test this, we analysed the relative proportion of ISC and EBs in various arrangements in the anterior midgut and found the majority of EBs do exist in ISC/EB pairs. However, we also found that a small proportion of EBs exist in isolation, which may indicate self-regulation independent of parental ISC. During the preparation of this manuscript, a live imaging study has also shown that the EB is a stable cell type distinct from the parental stem cell (i.e. not only as a transient progenitor) (Marchetti et al., 2022). In support of this hypothesis, our time-lapse imaging data have shown that EB cells indeed have motile behaviour.

In order to coordinate migration, EBs must be able to sense their environment. We believe EB responses are highly dynamic and are dependent upon the extent of tissue damage. While quiescent EBs often only contain small protrusions (Figure 4I’), activated EBs found in “regenerative” midguts can possess extensive protrusions that extend into the intercellular space between nearby ECs (Figure 6F, G, arrowhead). Previously, EB protrusions have only been observed in non-fixed tissues (Antonello et al., 2015a; Rojas Villa et al., 2019). We speculate that these protrusions are used for movement as well as signal detection, since cell protrusions have long been described to be essential for cell-cell communications (Buszczak et al., 2016). Such protrusions may only be transient and associated with early EB activation (Rojas Villa et al., 2019), hence not more commonly reported.

EBs also responded to DSS-induced damage. Consistent with our expectations, we found a significant increase in the cytoplasmic movements of EBs in the DSS-treated group compared to control, and this motile behaviour was abolished when we disrupted the actin cytoskeleton with *Rac1* RNAi. However, it is worth noting that there is a caveat associated with using *Rac1*.RNAi to disrupt the actin cytoskeleton. In mammalian studies, it has been shown that Rac1 not only controls the actin cytoskeleton but can also mediate the production of ROS (Ellenbroek and Collard, 2007). Elevated ROS levels are essential to damage-induced ISC proliferation in *Drosophila* (Buchon et al., 2009a). In the adult midgut, loss of Rac1 in progenitor cells can suppress ISC proliferation in the face of *Pe* infection (Myant et al., 2013). Therefore, we cannot discount the possibility that the *Rac1*.RNAi phenotype is somehow associated with the reduction in the number EBs. The active formation of EB clusters was also observed in the DSS-treated group, where EBs in close proximity extended their lamellipodia and came together. This behaviour appeared directional. Interestingly, the increased EB motility when subjected to DSS was only observed in cytoplasmic displacement but not nuclei displacement measured through trackmate. We propose the lack of significant nuclei displacement to be due to the short duration of our time-lapse imaging. The cytoplasmic displacement is a measure for protrusion formation and retractions, a rapid process as seen in ISCs (Hu et al., 2021). Hence, although we were able to capture EB protrusion dynamics within our 4-hours imaging window, perhaps nuclei displacement is a comparatively slower process which would require longer-term tracking of EBs after damage to be captured.

From our data, we know that single EB MET can happen, thus the question arises as to how cooperative MET occurs. One possibility is that EB cooperative behaviour is directly linked to the level of cell death (i.e. the number of ECs that require replacement) (Figure 10A). Here we propose a working model underlying EB clusters formation and midgut regeneration: EB activation and directional migration is triggered by the extent of local EC damage, whereby the threshold signal may be concentration dependent (for secreted ligands) or mechanical in nature. The initial gain of EB motility can be considered as a partial EMT event. Subsequently, the aggregation of EBs forms a hub for tissue repair that further recruits nearby EBs in a snowballing fashion. As a result, creating regions of high and low EB density reflected by our F-function measurements. At the site of damage, cooperative MET takes place to replenish lost ECs through EB-to-EC terminal differentiation. The new ECs generated based on this model would form the pattern of a localised cluster instead of being in isolation. Finally, the members of the regenerative cluster that did not undergo terminal differentiation disperse, thus the midgut tissue returns to homeostasis (Figure 10B). The snowballing mechanism proposed is based on our findings as well data inferred from two previous publications on *Sox21a* mutants. When Sox21a is mutated, EB differentiation is blocked, leading to an accumulation of undifferentiated EBs tumours. These tumours were also found to release Upd2, a cytokine sufficient to induce ISC proliferation (Jiang et al., 2009; Zhai et al., 2015). Therefore, we propose that the active aggregation of EBs could also stimulate further EB production, thereby producing a positive feedback loop.

**Figure 10.**
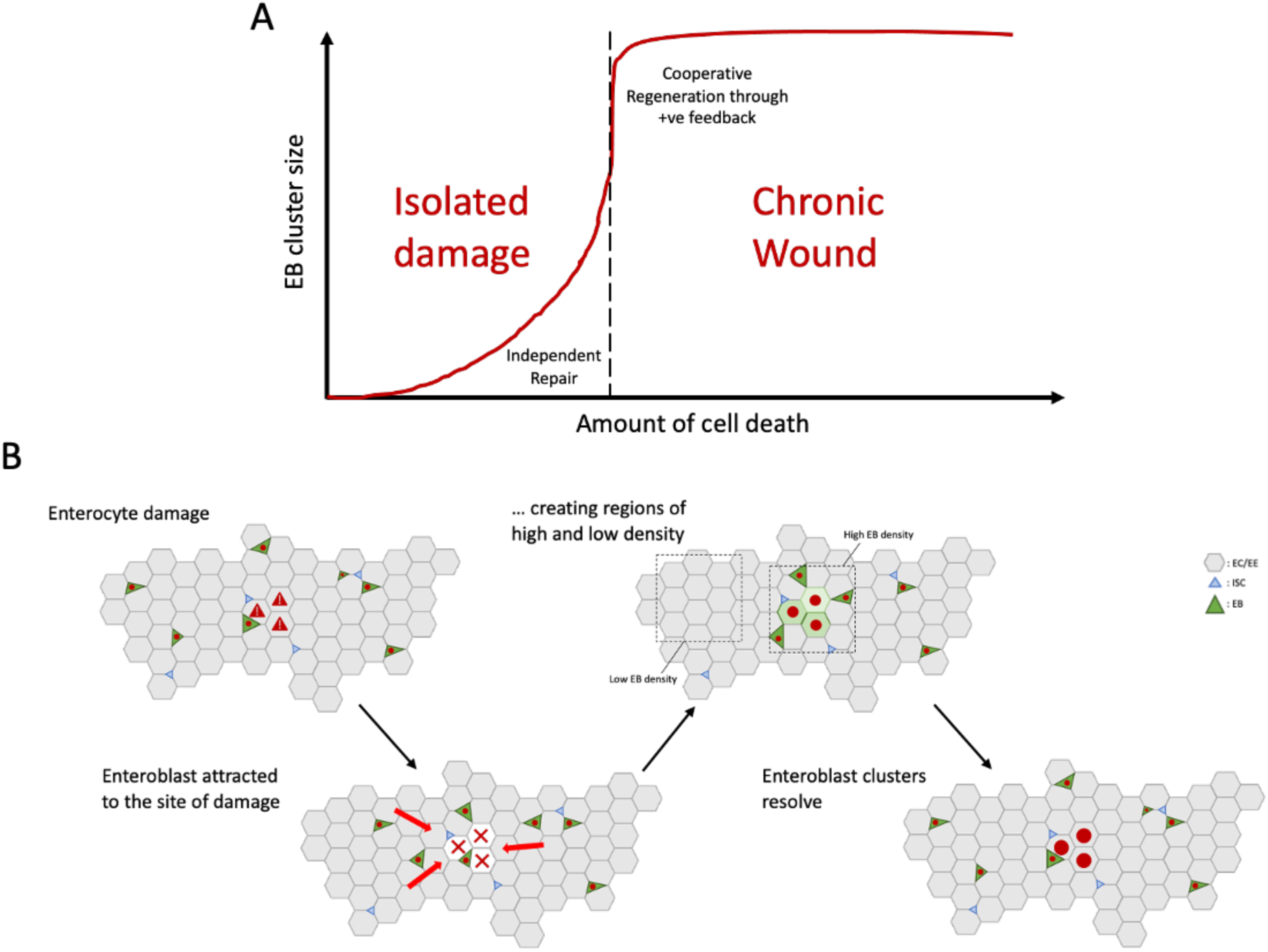
Model – Enteroblasts are required for site-directed regeneration. (A) EB-dependent adult midgut repair mechanisms are correlated with the extent of damage (i.e. amount of cell death). In the face of isolated damage, independent repair is likely to be facilitated through single cell MET. However, when the wound size (i.e. number of EC lost) reaches a threshold level, this could induce a cooperative MET process where multiple EBs are recruited to the damage site and facilitate tissue regeneration as demonstrated in panel B. (B) Model for EB directed migration to the damage site and the regeneration of ECs. In the presence of EC damage, nearby EBs are attracted to and migrate towards the site of damage through unknown mechanisms (possible mechanisms include: chemotaxis and/or mechanotransduction), forming “MET regenerative clusters”. The active migration of ‘stock’ EBs creates regions of high and low EB density within the midgut. Next, EB MET occurs and replenishes lost ECs. Finally, the EB clusters resolve and the tissue returns to homeostasis.

We then utilised the regenerative clusters to screen for candidate genes mediating EB response to tissue damage. If EB migration is the underlying force behind cluster formation, disruption of any component that regulates cell migration should be expected to produce a phenotype. The validity of this experimental set up was affirmed with identifying the GPCR Frizzled 2 (*fz2*), a canonical Wnt signalling ligand, in the cluster analysis. The canonical Wnt signalling is mainly involved in cell fate decisions, and is required for the midgut regenerative response to bacterial damage (Cordero et al., 2012). Recently, Hu et al. (2021), also described a role for Fz2/Dsh signalling in the migration and protrusion formation of ISCs, two components that are critical for regeneration (Hu et al., 2021). Here, we suggest that Fz2 possesses a similar role in EB migration.

It has been long established that mechanical cues can cause dynamic cytoskeleton rearrangements in cells and drive migration (Petzold and Gentleman, 2021) though studies on the function of mechanosensors (voltage-gated ion channels) in epithelial maintenance is currently limited. However, in the adult midgut, two recent publications have shown a role for mechanosensors in ISCs and EEs (Gong et al., 2022; He et al., 2018) strengthening the likelihood that EBs possess the same ability.

We have successfully identified a few mechanosensors and surface receptors for chemotaxis that warrant further research. This proof-of-concept screen highlights the power of a more global approach in studying gene function within a biological system, through coupling genetic manipulations with morphological analyses.

Focal adhesions (FA) function to anchor the cytoskeletal contractile machinery of a cell to the extracellular matrix, hence providing traction for the cell to move forward (Geiger et al., 2009). Unsurprisingly, EB-specific knockdown of Pax, a FA docking protein that regulates the Rho GTPases (Winkelman et al., 2020), significantly reduced regenerative clusters formation. For the occasional clusters that formed, they were made of very few EBs. Activation of Rho induces FA formation, and subsequently lamellipodia (Chen et al., 2005), both essential for cell migration.

RNAi knockdown of several mechanosensors had presented novel functions in that they either disrupted EB regenerative cluster formation only (*nanchung*, *cacophony*), or as well as EB terminal differentiation (*sandman*). *Sandman* (*sand*) encodes a K2P channel subunit known to buffer K^+^ levels in the cortex glia (Weiss et al., 2019), and its mammalian ortholog TREK-1 regulates the influx of K^+^ current in response to membrane stretching, thus induces actin cytoskeleton remodelling and the formation of membrane protrusions (Lauritzen et al., 2005). Therefore, we can infer a similar function for Sand in EB migration. Nanchung (Nan) is a member of the TRP channel family and its mammalian ortholog TRPV5 is activated by mechanical stress past a certain threshold (Shen et al., 2015). Finally, *cacophony* (*cac*) encodes a voltage-gated calcium channel subunit (Kawasaki et al., 2002), the induction of Ca2^+^ pulses at the leading edge of a cell had been shown to direct cell migration (Tsai et al., 2015). Since most TRP channels mediate Ca^2+^ entry (Gees et al., 2010), it is feasible that TRP channel components control EB cell migration through mechanotransduction.

Switching to the cell surface receptors, many diffusible ligands are known to mediate proliferative responses in the midgut, so it is entirely possible that they also guide EB migration via chemotaxis (SenGupta et al., 2021). Here we briefly discuss two candidates that, when knocked down, led to the most significant alteration to the EB clusters: *fra* and *dome*. Frazzled is a Deleted in Colorectal Carcinoma (DCC)-family receptor that mediates axon guidance and embryonic midgut formation in *Drosophila* (Pert et al., 2015; Tessier-Lavigne and Goodman, 1996). We propose Fra can mediate EB migration and formation of regenerative clusters in a similar fashion. Interestingly, though *fra* is expressed in EBs, its ligands netrin-A (*netA*) and netrin-B (*netB*) have minimum expression levels in all cell types of the midgut (Li et al., 2022). This may be explained by a line of evidence from our previous work that in embryonic mutants for *netAB^Δ^*, which deletes both *netA* and *netB*, midgut cells were still able to form protrusions and exhibit patches of non-polarised F-Actin (Pert et al., 2015). This suggests Frazzled can function in a netrin-independent manner. Alternative ligands for Fra need to be considered for EBs to respond to a morphogen gradient, such as Nolo, an ortholog of MADD-4 which physically binds to UNC-40 (a Fra ortholog) in *C. elegans* (Seetharaman et al., 2011).

Domeless is the receptor for JAK/STAT signalling and is highly expressed in both the EB and late EB clusters, though not ISCs (Li et al., 2022). This suggests that Dome possesses a distinct function in EBs. We know that ISC proliferation in response to environmental challenge is mediated through JAK/STAT signalling in EBs (Zhou et al., 2013), and JAK/STAT activation in EBs is essential for their differentiation into ECs (Beebe et al., 2010). We expect the downstream mechanism of Dome in regulating EB migration to be different from that of ISC regulation and terminal differentiation, and may be able to draw parallel to the known regulatory function of Dome in border cell migration during oogenesis (Ghiglione et al., 2002). Together, we have identified several potential regulators of EB migration and MET that warrant further follow up. More targeted experiments are required to determine the mechanism of function for these genes identified, how exactly do they impact EB regenerative cluster formation.

There are still a number of open questions yet to be addressed: 1. How far can EBs move (i.e. is there a “regenerative domain” supported by individual EBs, as there is with ISCs)? 2. Why is there a large variation in EB cluster size? Perhaps it is associated with the extent of tissue damage. And 3. Do EBs form a regenerative cluster through directional migration? To verify this, a more precise and repeatable method of damage will be required in conjunction with live imaging. Regardless of the precise mechanisms required, we have established that EBs are an active participant in MET regenerative cluster formation and are therefore involved in regeneration.

In summary, our results demonstrated the ability to extract meaningful biological data out of currently underutilised image data. Using the morphometric and distribution analysis pipeline, we uncovered a previously unreported bimodal distribution pattern of EB cells with a functional role in tissue homeostasis and regeneration. We proposed a new model where EB-directed tissue regeneration is a collective and coordinated process, and using the presence of MET regenerative clusters, identified several novel regulators. It will be important now to confirm EB directional motility and have a localised inducible damaging system for more in-depth analysis of candidate gene functions.

## Movies

Movie 1: Representative movie of midgut tissue health post-imaging. Peristaltic contractions are visible indicative of normal tissue function.

Movie 2: Representative movie of wildtype EB behaviour. Limited motility is observed over a 4-hours imaging window.

Movie 3: Representative movie of EB behaviour when the midgut is damaged through DSS-feeding. Formation of extended membrane protrusions and cell displacements are observed over a 4-hours imaging window.

Movie 4: Representative movie of EB behaviour with Rac1 RNAi knockdown. Cells appear more rounded with limited motility over a 4-hours imaging window.

Movie 5: Representative movie of Rac1.IR EB behaviour when the midgut is damaged through DSS-feeding. Membrane protrusions are still visible though appear restricted in large scale movements over a 4-hours imaging window. One cell cluster with extended protrusion and cell displacement is seen.

## Acknowledgments

We thank: Maria Dominguez for the kind gift of the ReDDM fly stocks, and Mikio Furuse for the anti-Ssk and anti-Mesh antibodies; The Bloomington Drosophila Stock Centre, DGRC and the Vienna Drosophila Research Centre for fly stocks, and the Developmental Studies Hybridoma Bank for antibodies; The University of Melbourne Biological Optical Microscopy Platform, and the Monash University OzDros fly importation service; Gary Hime for helpful reading of the manuscript. This work was supported by a Melbourne Research Scholarship to F.Z. and a University of Melbourne MRGSS grant and National Health and Medical Research Council Project Grant (GNT1107123) to M.J.M.

